# Genomic evolution of pancreatic cancer at single-cell resolution

**DOI:** 10.1101/2024.08.03.605165

**Authors:** Haochen Zhang, Palash Sashittal, Elias-Ramzey Karnoub, Akhil Jakatdar, Shigeaki Umeda, Jungeui Hong, Anne Marie Noronha, Agustin Cardenas, Amanda Erakky, Caitlin A. McIntyre, Akimasa Hayashi, Nicolas Lecomte, Wungki Park, Nan Pang, Eileen M. O’Reilly, Alice C. Wei, Benjamin J Raphael, Christine A. Iacobuzio-Donahue

**Affiliations:** Gerstner Sloan Kettering Graduate School of Biomedical Sciences, Memorial Sloan Kettering Cancer Center, New York, NY, USA; Human Oncology and Pathogenesis Program, Memorial Sloan Kettering Cancer Center, New York, NY, USA; Department of Computer Science, Princeton University, Princeton, NJ, USA; David M. Rubenstein Center for Pancreatic Cancer Research, Memorial Sloan Kettering Cancer Center, New York, NY, USA; Marie-Josée & Henry R. Kravis Center for Molecular Oncology, Memorial Sloan Kettering Cancer Center, New York, NY USA; Division of Surgical Oncology and Endocrine Surgery, The University of Texas Health Science Center San Antonio, San Antonio, TX, USA; Department of Pathology, Kyorin University, Mitaka City, Tokyo, Japan; Department of Medicine, Memorial Sloan Kettering Cancer Center; Department of Surgery, Memorial Sloan Kettering Cancer Center, New York, NY, USA; Department of Medicine, Weill Cornell Medicine, New York, NY, USA; Department of Pathology and Laboratory Medicine, Memorial Sloan Kettering Cancer Center, New York, NY, USA

## Abstract

We adapted a previously developed targeted single-nucleus DNA sequencing (snDNA-seq) method and constructed a new suite of computational analysis tools to study 137,491 single-nucleus DNA libraries from 24 pancreatic cancers collected under a variety of clinical scenarios including early and late diagnoses, different metastatic sites and before- and after-treatment. We refined the mutational landscape of pancreatic cancer by capturing events missed by bulk sequencing, and validated the evolution pattern of early fixation of driver single-nucleotide variants (SNVs) followed by generation of intratumoral heterogeneity for copy number variations (CNVs). Intertumoral convergent evolution was common, including subclonal inactivation of TGF-β pathway by mutating various components of it; intratumoral convergence was rarely observed, likely due to strong selective force in pancreatic cancer development. Continuous evolution was frequently seen manifesting as CNVs. In the context of non-targeted treatments, no particular pattern was found across metastases or through treatment. In six pancreatic cancers with germline *BRCA2* mutation, we discovered varied timing of biallelic inactivation of *BRCA2,* which sculpted different evolutionary trajectories and could presumably contribute to differential response to treatment. As the first large-scale application of targeted snDNA-seq on pancreatic cancer, this work provides a sample processing and computational analysis pipeline that warrants further clinical utility.

## Introduction

Pancreatic ductal adenocarcinoma (PDAC) is one of the most lethal cancer types with a 5-year survival of less than 12%^1^. This rate has increased minimally over the past decades despite improvements in surgical and medical management, with PDAC projected to become the second most common cause of cancer-related deaths in the United States^2^. In recent years, collaborative research efforts across the globe have advanced our understanding of the disease’s molecular underpinning and revealed several opportunities for precision medicine. Novel therapeutic approaches such as PARP inhibitors, KRAS inhibitors, and immunotherapies are being developed to target individual tumors’ vulnerabilities with promising results^3–6^.

Numerous next-generation sequencing (NGS) studies have revealed PDAC clonal evolution as a hybrid model of stepwise and punctuated events. Point mutations in multiple driver genes (i.e. *KRAS, CDKN2A, TP53*) followed by allelic loss of the wild type alleles are gained through progressive waves of clonal selection^7^, while pervasive copy number variations or larger scale genomic rearrangements that further grant growth advantages are acquired rapidly following polyploidization. Polyploidization coincides temporally with loss of the wild type p53 allele and the acquisition of invasive potential^8,9^. Elucidation of these genomic events has largely been accomplished by sequencing of bulk tissues; in some cases resolution has been increased by use of microdissected samples and/or development of computational tools to deconvolve the complexity of subclonal mixtures^10–12^. Regardless, given clonal evolution happens at the single-cell level one must rely on a set of strong assumptions which may overlook important events such colocalization of alterations within a singular genome or mutual exclusivity indicating distinct populations. Thus, single-cell DNA sequencing is required to elucidate PDAC’s clonal evolution and heterogeneity above the highest resolution achieved by bulk sequencing, which currently requires a mutation to be present in approximately 2% of cancer cells^13^ (**Methods**).

We previously optimized a scalable workflow to apply high-throughput single-nucleus DNA sequencing (snDNA-seq) to archival PDAC samples^14^. Here we utilized this workflow, in combination with an accompanying set of novel computational methods and unique patient sample set to generate the highest resolution view of genomic evolution to date. In doing so, we aim to define at single cell resolution the genomic bottlenecks and intercellular heterogeneity present over the course of PDAC evolution spanning early invasion to dissemination to a variety of secondary sites.

## Results

### 1. PDAC mutational and evolutionary landscape at single-nucleus resolution

#### Cohort overview

We studied the single cell genomic features of 24 patients with a primary pancreas neoplasm. The cohort included samples from patients with both early-stage disease collected from surgical resections (n=10 patients) and late-stage unresectable disease from research autopsies (n=11 patients) (**Figure 1A, Supplementary Table 1)**. Matched longitudinal biopsies taken by 18/22-guage core needle biopsies before and after treatment were collected from 6 patients (**Supplementary Table 2)**. The final pathologic diagnosis for 23 patients was PDAC; 1 with acinar cell carcinoma (ACC). We opted to study the latter patient given their clinical sequencing result suggested a germline mutation in *BRCA2 (*gBRCA2); 5 of the PDAC patients had documented gBRCA2 mutations as well, providing an opportunity to understand genomic features of neoplasms that arise in the setting of this set of high-risk germline variants. Collectively, a total of 30 primary tumor samples and 42 metastases were studied. For 21 patients, at least two spatially distinct samples were studied per patient (range 2-9). Metastases from 35 secondary sites were sampled in the autopsy participants with a median of 2 metastases (range 1-7) studied per patient.

**Figure 1:**
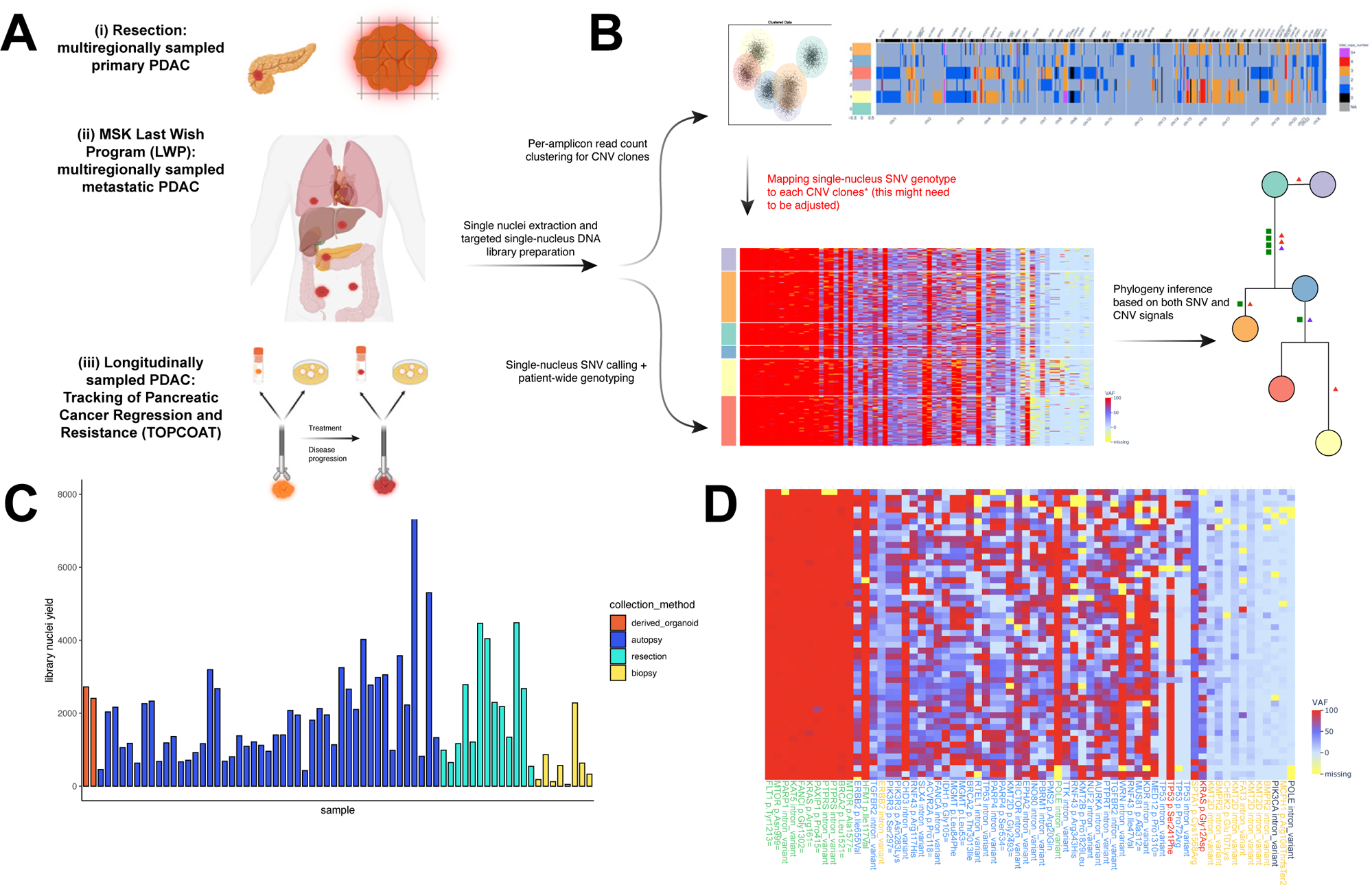
Sample collection, library preparation and computational analysis workflow. **a.** Sample collection procedures. **b.** Overview of computational analysis workflow. Top: copy number variation (CNV) clone clustering procedure and example result. Bottom left: single-cell mutation heatmap, where the heat represents the variant allele frequency (VAF) of each mutation in each single cell. Bottom right: example result phylogeny from fast-Constrained Dollo Reconstruction (fast-ConDoR). **c.** The number of nuclei collected for each sample’s library, colored by collection method. In total N=74 Tapestri snDNA libraries were constructed. **d.** Single-cell mutation heatmap of PC19 intervational radiology (IR) biopsy sample, which yielded the lowest library size of 50 single cells. Variants are labeled based on comparison with bulk sequencing and with panel of normal (PoN) and cohort-wide blacklist. Green: germline homozygous variants. Blue: germline heterozygous variants. Red: somatic variants detected in bulk sequencing. Yellow: likely artifact variants. Black: others.

#### Genetic Features of Cohort

To discern the genetic features of our cohort, we performed targeted snDNA-seq using a custom designed 596-amplicon panel targeting essential genes of primary pancreatic neoplasms; genes affected by both single nucleotide variant (SNV), small insertion/deletion (indel) and/or copy number variations (CNV) were considered (**Supplementary Table 3; Methods**). We custom designed a bioinformatics and computational analysis pipeline for this snDNA-seq dataset (**Figure 1B; Supplementary Figure 1A, B**).

In total 137,491 single-nucleus DNA libraries were profiled by this panel with a mean of 5,729 per patient (range 329 – 20,123) and 1,858 single nuclei studied per sample (range 50-7,540) (**Figure 1C**). Bulk whole-genome/exome sequencing (WGS/WES) data were collected for each tumor and matched normal sample simultaneously for quality control of germline/somatic event calling.

While resected/autopsied samples were large enough to be embedded in optimal cutting temperature (OCT) compound and inspected to ensure cellularity and tumor purity, longitudinal biopsies were usually too small (<8mm^3^) to be embedded. As a result, these biopsies were extracted and sequenced without controlling for the two variables, resulting in many such samples with lower nuclei yields. Nevertheless, even those biopsies with the lowest yields (i.e. PC18) resulted in useful genotype information (**Figure 1D**), thus proving the feasibility of applying this snDNA-seq approach to routine clinical biopsies.

We detected somatic SNVs, indels, monoallelic and/or biallelic losses in at least one known driver gene in all cases analyzed (**Figure 2A**). Activating *KRAS* mutations were identified in all but two PDAC (21of 23, 91%). Targeted clinical sequencing performed for these two neoplasms revealed a *BRAF* fusion and an NRG fusion, respectively, which were not on our panel. No *KRAS* mutation was found in the ACC. Alterations in *TP53* (75%), *CDKN2A* (71%), and *SMAD4* (71%) were also observed at high frequency, as expected^15^. Somatic alterations noted at lower frequency included *SMAD2, SMAD3, TGFBR1, TGFBR2, ARID1A, ARID2, BRCA2* and *RNF43.* Copy number alterations identified included amplifications in *MYC* (46%)*, GATA6* (17%) and *MTOR* (4%, 1 case), as well as biallelic loss (homozygous deletion, “homdel”) in genes such as *CDKN2A* (67%)*, SMAD4* (25%) and *TGFBR2* (8%, 2 cases). A technical strength of snDNA-seq compared to bulk sequencing is that it enables direct calculation of each alteration’s corresponding cancer cell fraction (CCF) (**Methods**); as a result, we could distinguish between clonal drivers such as *TP53* (mean CCF = 0.90), *BRCA2* (0.88), *CDKN2A* (0.83), *MYC* (0.83), and subclonal drivers such as *SMAD4* (0.68), *TGFBR2* (0.66), *ARID1A* (0.51) (**Figure 2A, square size**).

**Figure 2:**
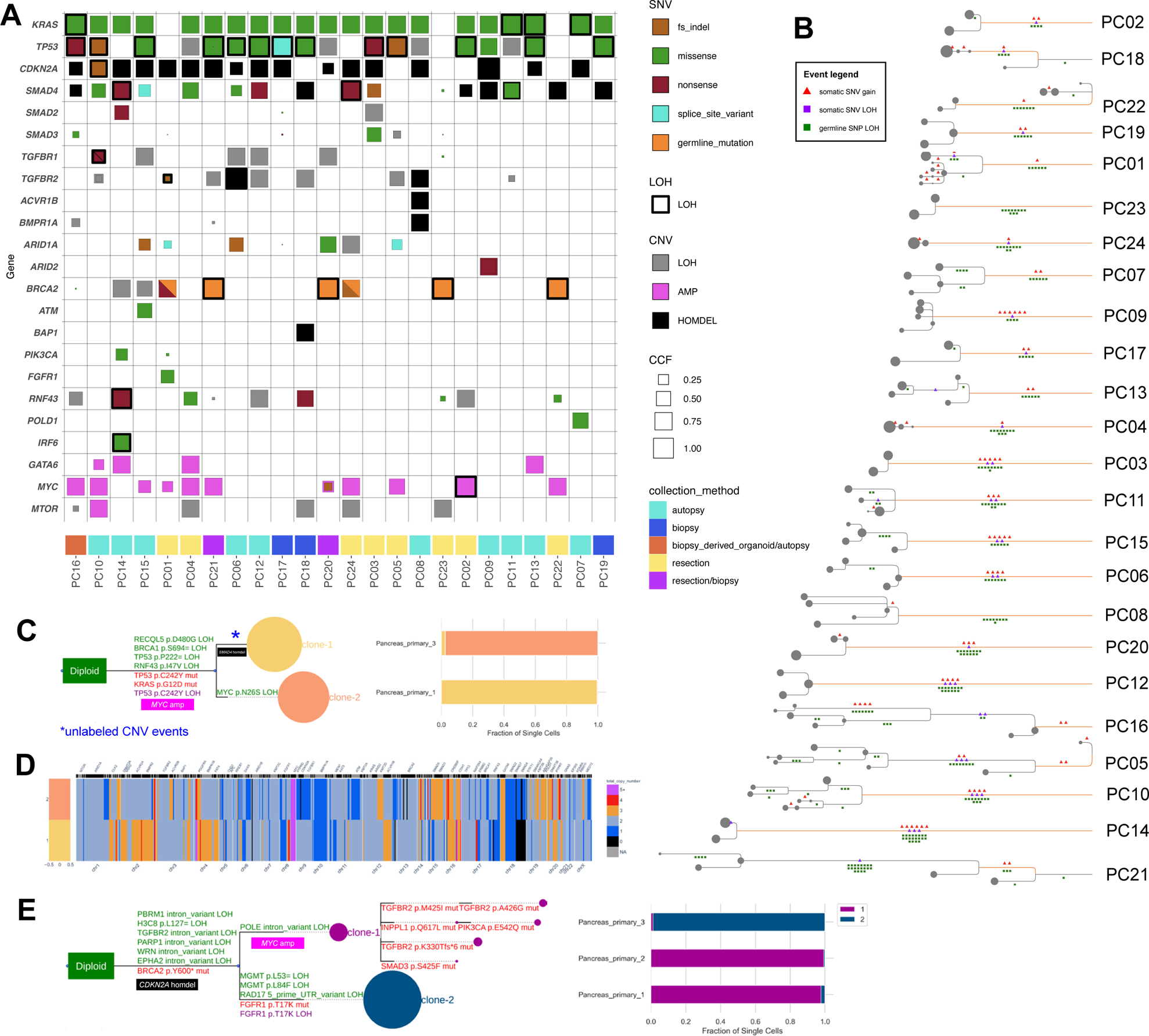
Refined mutational landscape, evolutionary patterns of pancreatic cancer revealed by snDNA-seq. **a.** Oncoprint depicting the type of mutations and their cancer cell fraction (CCF) in each pancreatic cancer case studied. On the x-axis: cases are sorted by their respective number of mutational events in descending order from left to right. On the y-axis: genes are sorted by the number of alterations in the patient cohort in descending order from top to bottom. Within the grid, the square size is proportional to the CCF of each alteration. The square color corresponds to the alteration type, which is of two categories: single-nucleotide variants (SNVs) and copy number variations (CNVs). The thickened border represents loss of heterozygosity (LOH) status. **b.** Overview of single-cell clonal phylogenies inferred for each case. The branch length on the x-axis represents inferred phylogenetic distance among clones (grey circles). The orange branch is the inferred trunk of the phylogeny (Methods). The size of the circle is proportional to the number of cells in that clone. For events on the edge: red triangles 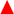 indicate gain of somatic SNVs; purple triangles 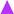 indicate LOH of somatic SNVs; green squares 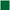 indicate LOH of germline SNPs. **c.** Phylogeny (left) and sample clonal proportion (right) of PC02. In the phylogeny, red texts indicate gain of somatic SNVs; purple text indicate LOH of somatic SNVs; green texts indicate LOH of germline SNPs; total copy number events are not labeled. **d.** Clone total copy number profiles of PC02. On the y-axis, clone colors correspond to those in c. The x-axis is the genomic coordinate, with chromosome numbers labeled at the bottom, gene names on the top. The heat represents the total copy number state of each clone at each genomic location. **e.** Phylogeny (left) and sample clonal proportion (right) of PC01.

The snDNA-seq approach expands point mutation (SNV/indel) calling *sensitivity* from bulk sequencing’s 2% to 0.05% single-cell prevalence, approximately. Yet we estimated its *specificity* to be limited to 0.5% single-cell prevalence (**Methods**). At this threshold, we observed a subset of genes with an increased frequency of point mutations than reported in bulk sequencing studies. For example, *ARID1A* mutations were present in 29% of our cohort compared to 6% in The Cancer Genome Atlas (TCGA) (chi-squared test, p=0.00015) and 14% in the International Cancer Genome Consortium (ICGC) (p=0.075)^16,17^. Furthermore, subclonal and subgene (affecting only a part of the gene) monoallelic/biallelic losses are difficult to detect with common bulk sequencing methods, particularly for PDAC where tumor purity is often limited. However, our targeted snDNA-seq method evaluated each mutant loci’s variant allele frequency (VAF) and single amplicons’ (200-300 base pairs) read count while accounting for amplicon dropout rates, to detect such events in small groups of cells (**Methods)**. For instance, we could visualize homdel confined to a single amplicon of *SMAD4* in the neoplastic clone of PC18 (**Supplementary Figure 1C)**. Across the cohort we detected a higher alteration (including both short mutation and monoallelic/biallelic loss) frequency of *CDKN2A* and *SMAD4* (71% and 71%, respectively) than reported by TCGA (30%, 32%) and ICGC (30%, 27%). Specifically, we detected homdel of *CDKN2A* in 67% cases *and SMAD4* in 25% cases, which are comparable to the numbers (around 60% for *CDKN2A* and 40% for *SMAD4*) found in a mixed cohort of primaries/metastases using microdissection to enrich for tumor content^11^. We also detected a high frequency of monoallelic losses (loss of heterozygosity, LOH) affecting *TGFBR1* (21%) and *TGFBR2* (29%). We interpret these findings to reflect the increase in sensitivity by our assay rather than more extensive sampling, as these rates are also higher than reported in a recent study of 91 multiregionally sampled PDAC research autopsies (work in review).

#### Major patterns of evolutionary trajectory

In addition to novel methods of calling somatic variants, we developed fast Constrained Dollo Reconstruction (fast-ConDoR), a new computational approach to reconstruct the single-cell clonal phylogeny from each patient. Briefly, this method first clusters cells into CNV clones, and uses germline single-nucleotide polymorphism (SNP) and SNV allelic information to build a clonal phylogeny at single-cell resolution^18^ (**Figure 1B**, **Methods)**.

**Figures 2B** illustrates all 24 phylogenies generated from this method. A mean of 2.63 truncal driver SNV events (**Methods)** were covered by our panel, consistent with the number calculated unbiasedly in a work in review; among all driver SNVs identified, on average 74.4% were truncal in origin (i.e. occurred before the most recent common ancestor (MRCA) of all clones identified). This informed us that the major distinction among PDAC clones CNVs, as illustrated by the example case in **Figure 2C-D**: most of the SNV events and relevant LOHs were on the trunk, while total copy numbers varied across the genome between the two clones. This pattern of early fixation of drivers followed by generation of intratumoral heterogeneity by CNVs is consistent with multiregional bulk sequencing of larger PDAC cohorts^19^. The number of subclones resolved per case was not correlated with the number of single nuclei analyzed, as demonstrated by two edge cases: PC11 had one of the highest patient-level library size (16,139 single nuclei) but only 3 tumor clones, while PC21 had only 329 single nuclei but 4 tumor clones.

#### Evolutionary patterns of PDACs with germline BRCA2 alterations

While patients with germline mutations in the *BRCA2* gene are associated with an inherited risk of developing PDAC as well as increased sensitivity to platinum salts, the evolutionary features of how pancreatic neoplasms arise in the setting of gBRCA2 is poorly understood. We therefore studied 6 such cases in our cohort, all of whom had heterozygous gBRCA2 mutations and biallelic *BRCA2* inactivation through different mechanisms. PC20, PC21, PC22, PC23’s tumor cells lost the wild-type *BRCA2* allele (LOH) (**Supplementary Figure 2A**), while PC01 and PC24 had inactivating somatic mutations to *BRCA2*. Mutation signature analysis confirmed the presence of the HRD phenotype in all 6 cases (**Supplementary Figure 2B)**. PC05, a non-gBRCA2 carrier initially intended as a control, had prevalent HRD signature as scored by HRDetect **(Methods)**. This patient was noted to have an elevated familial cancer propensity, potentially linking the HRD phenotype with unknown germline mutation(s).

The mean CCF for BRCA2 alteration was 0.88 (**Figure 2A)** suggesting generally early inactivation of the gene. Nevertheless, review of phylogenies revealed heterogeneity with respect to this timing. For example, PC01’s phylogeny showed that the somatic *BRCA2* nonsense mutation was truncal, followed by several other LOH events and *CDKN2A* homozygous deletion (**Figure 2E, left)**. The phylogeny subsequently diverged into two mutually exclusive subclones carrying an *FGFR1* p.T17K mutation with LOH, or a *TGFBR2* p.K330Tfs*6 mutation with LOH. The subclones separated spatially (**Figure 2E, right).** WGS performed on the same tissue identified a *TRIM24-BRAF* translocation, a *KRAS-*independent route towards oncogenic MAPK activation, suggesting that this PDAC developed through noncanonical mechanisms. Similarly, case PC23 showed a truncal *BRCA2* LOH **(Supplementary Figure 2C)**, with no detectable alteration to other canonical PDAC driver genes; WGS suggested an *NRG* fusion, another noncanonical oncogenic mechanism converging to the same pathway as *KRAS*.

By contrast, in case PC20 the WT *BRCA2* allele was likely lost later in the evolutionary life history of the neoplasm. Although technical allelic dropout at the *BRCA2* loci made it challenging for single-cell genotyping to accurately time the occurrence of *BRCA2* LOH in relation to other common drivers, in this case we observed a small population of cells containing a heterozygous germline *BRCA2* p.I2627F mutation, a heterozygous somatic *KRAS* p.Q61H mutation, a *CDKN2A* homozygous deletion and a homozygous somatic *ARID1A* p.R1989P mutation; LOH of at least 8 additional germline SNPs were also noted (**Supplementary Figure 2D**). The *ARID1A* p.R1989P homozygous genotype and germline SNPs’ LOH validated this population as non-doublet and indicated that *BRCA2* LOH likely came after the driver SNV and CNV events. In PC21, although we didn’t observe a subclone with the genotype as PC20’s, we observed a tumor subclone where *BRCA2* was homozygous deleted, in contrast to the main tumor population where *BRCA2* had LOH (**Supplementary Figure 2E, F)**. Both populations had the canonical drivers *KRAS* p.G12D and *TP53* p.L344Q. However, orthogonal statistical testing on raw single-cell read count data validated that the *BRCA2* homdel significantly colocalized with a heterozygous *TP53* state (p=0.0078), substantiating the distinct subclonal population. This also suggested a late selective pressure for *BRCA2* inactivation.

In PC22 the timing of loss of the WT BRCA allele could not be determined as the non-tumor population also showed elevated allelic imbalance (**Supplementary Figure 2A)**. PC24’s somatic *BRCA2* p. p.S1832Ifs*2 mutation was not covered by our snDNA-seq panel; yet WGS suggested a lower CCF (0.80) as compared to the two main drivers *KRAS* p.G12D (0.85) and *SMAD4* p.Q116* (1.00), categorizing it into the late-BRCA2 biallelic inactivation group.

In sum, we found that while *BRCA2* biallelic inactivation often takes place late in PDAC development and follows canonical drivers such as *KRAS, TP53, SMAD4* alteration, exceptions are present, where *BRCA2* took place early and could potentially drive PDAC in non-canonical mechanisms.

### 2. Convergent evolution towards TGF-β inactivation

Based on bulk sequencing one or more pathways in PDAC may be affected by convergent evolution. For example, *KRAS* wildtype (WT) PDACs have been shown to have alternative genetic mechanisms of activating MAPK-ERK signaling^16^, evident in our two *KRAS* WT cases: PC01 had *BRAF* fusion while PC23 had *NRG* fusion. The TGF-β pathway may accumulate genetic alterations in intracellular pathway components (i.e. *SMAD4, SMAD2*) or surface receptors (i.e. *TGFBR2*)^20^.

With snDNA-seq’s increased resolution, we asked if within the same tumor, additional clones exist with somatic mutations in these or other PDAC-relevant molecular pathways that have been unrecognized by bulk sequencing methods; this is important to know for efforts to manage resistant disease that can arise from pre-existent cells^21^. Single cell populations of >= 1% CCF were included in the oncoplot in **Figure 2A**, and showed mutations missed by bulk-sequencing but captured by snDNA-seq: *PIK3CA* mutation (CCF=0.0375), a *SMAD3* mutation (CCF=0.0134) in M04 and a *SMAD4* mutation in TP11 (CCF=0.0113). Nonetheless, even at this resolution we found that most PDAC driver pathways contain a single genetic event indicating prior strong selection in association with a near total to complete clonal sweep. The only exception was PC01, where three mutually exclusive clonal lineages carrying (1) *TGFBR2* p.K300Tfs*6 (2) *TGFBR2* p.A426G/p.M425I (3) *SMAD3* p.S425F mutations (**Figure 2E**) showed convergence towards TGF-β inactivation; in the same tumor, two mutually exclusively lineages carried (1) *FGFR1* p.T17K (2) *PIK3CA* p.E542Q respectively, demonstrating convergence towards MAPK–ERK activation.

As we extended the threshold to 0.5% single-cell prevalence (not necessarily limited to the cancer cell fraction) per sample, low-prevalence mutations to other PDAC drivers were detected but did not show strong positive selection (dN/dS = 1.04) **(Figure 3A, Methods)**. In fact, possible negative selection was observed when considering for all genes on the panel (dN/dS = 0.89), suggesting a purifying effect of cancer growth or the presence of these mutations in non-neoplastic cells. One secondary *KRAS* oncogenic alteration was identified in one section of the primary site of patient PC10 (single-cell prevalence = 0.0083) but showed little correlation with the main tumor clone (**Figure 3B)**.

**Figure 3:**
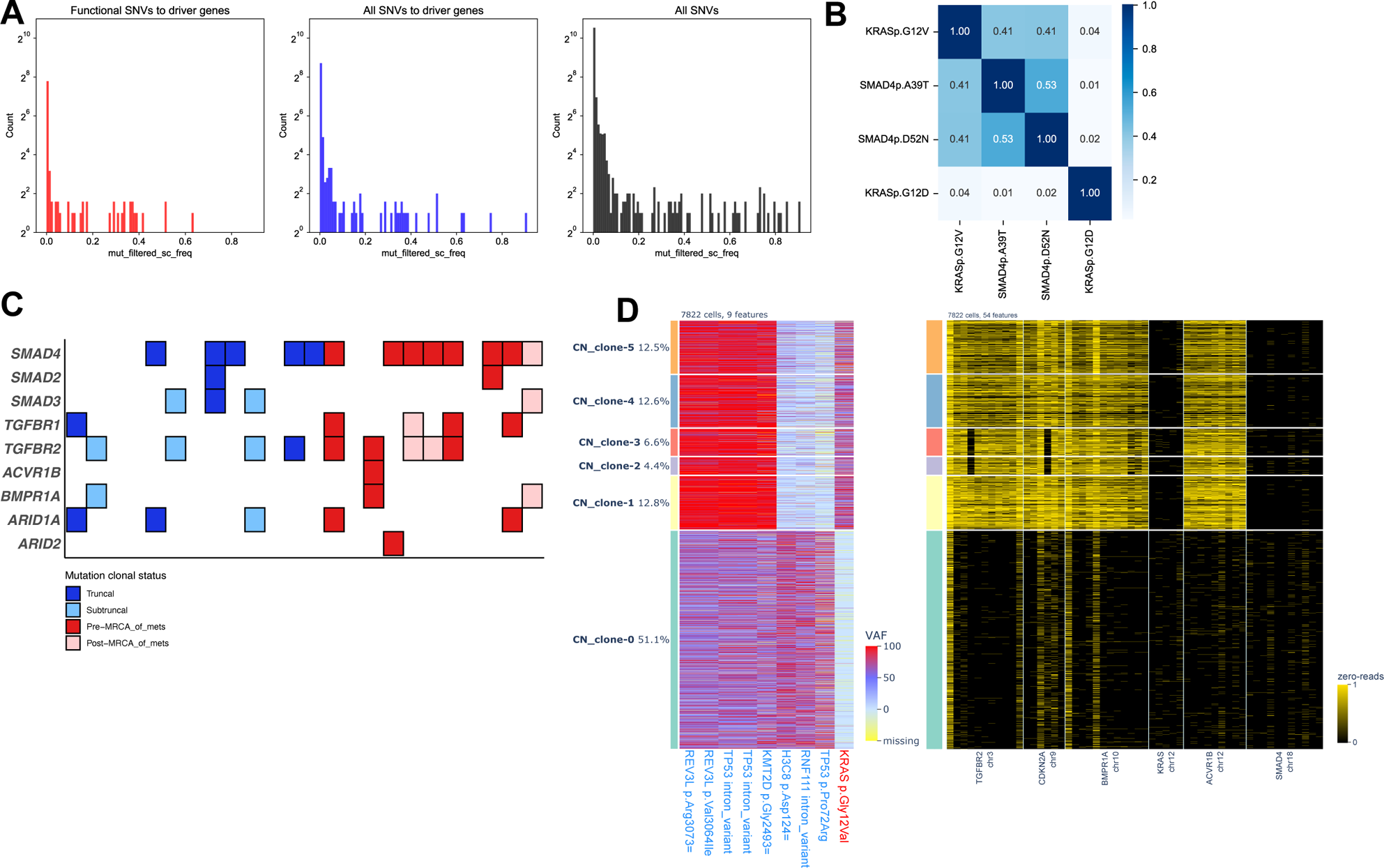
Convergent evolution in pancreatic cancer. **a.** Distribution of SNVs across varying mutational frequencies. X-axis represents within each case studied, the prevalence of single cells mutated for each SNV. This is not normalizing for tumor purity which means that a clonal driver like a KRAS mutation with CCF=1 might have 0.4 prevalence because of inclusion of 60% normal cells. Y-axis is the count of mutations. Left panel: distribution of functional SNVs to driver genes (Methods). Middle: all SNVs to driver genes. Right: all SNVs. **b.** In PC11, pair-wise statistical test for mutual exclusivity for SNVs of interest. The values are p-values so the lower the value, the more likely the pair of mutations are localizing in different cells. **c.** TGF-β mutations’ clonal status across all cases studied. For early-stage cases without metastasis, truncal vs subtruncal status is labeled; for late-stage cases with metastases, whether or not the mutations came before the most recent common ancestor (MRCA) of the metastatic clones is labeled. **d.** paired single-cell mutation heatmap and single-cell amplicon heatmap of mutations and genes of interest in PC08.

Unlike these isolated examples, we again found pervasive inter-patient convergence towards the phenotype of tumor cell-intrinsic TGF-β unresponsiveness (**Figure 3C**). The most common form of genetic inactivation of the pathway remains *SMAD4*, yet 6 other genes also have inactivating mutations that do or do not co-occur with *SMAD4*. We found that these alterations are subtruncal in many early-stage cases, while mostly occurring before the MRCA of metastases in late-stage cases. This suggests two features: (1) inactivating mutations of the TGF-β pathway have a strong selective advantage in the desmoplastic, nutrient-poor pancreas microenvironment; (2) these inactivating mutations have a selective advantage for survival in secondary sites. Some cases inactivated multiple components of the TGF-β pathway, but not *SMAD4*, the most prominent example being PC08, which had three TGF-β family receptors focally deleted (**Figure 3D**). We also noted examples of inter-patient convergence for inactivation of the SWI/SNF pathway. For example, 7 of 24 cases had an *ARID1A* mutation, and in one PDAC that was WT for *ARID1A* (PC09) we found two nonsense mutations in *ARID2* (**Figure 2A)**.

### 3. Continuous evolution driven by CNVs

In addition to convergent evolution, we noted several examples where somatic alterations continued to accumulate within a single gene and the same lineage over the course of PDAC evolution. For instance, in case PC06, *TGFBR2*’s second amplicon was deleted in virtually all cancer cells (**Supplementary Figure 3A**) whereas *CDKN2A* amplicons #1, 2, 6 were likely hemizygously deleted in clones 3 and 4 (**Supplementary Figure 3B, arrow-pointed)**; subsequently the second allele was lost which were only localized to clones 1 and 2 corresponding to the left liver metastasis (**Supplementary Figure 3C)**. Case PC10 (described as PA04 in our previous work^14^) had two independent mutations to *SMAD4* on the same chromosome, which translate to 13 codons apart, and then loss of the wildtype allele (**Figure 4A, left**). Homozygous deletion to 10 of 11 amplicons of *SMAD4* took place in the MRCA of all tumor clones (**Figure 4A, right**). The only intact amplicon was the one where the two point mutations located to. One subclone exclusively present in the liver metastasis and one region of the primary (**Figure 4B, C**) had a *TGFBR2* LOH event, which potentially further contributed to the phenotype of TGF-β inactivation^22^. This pattern was also observed in PC21, where the *BRCA2* locus with a germline mutation underwent both LOH and homozygous deletion **(Supplementary Figure 2E**).

**Figure 4:**
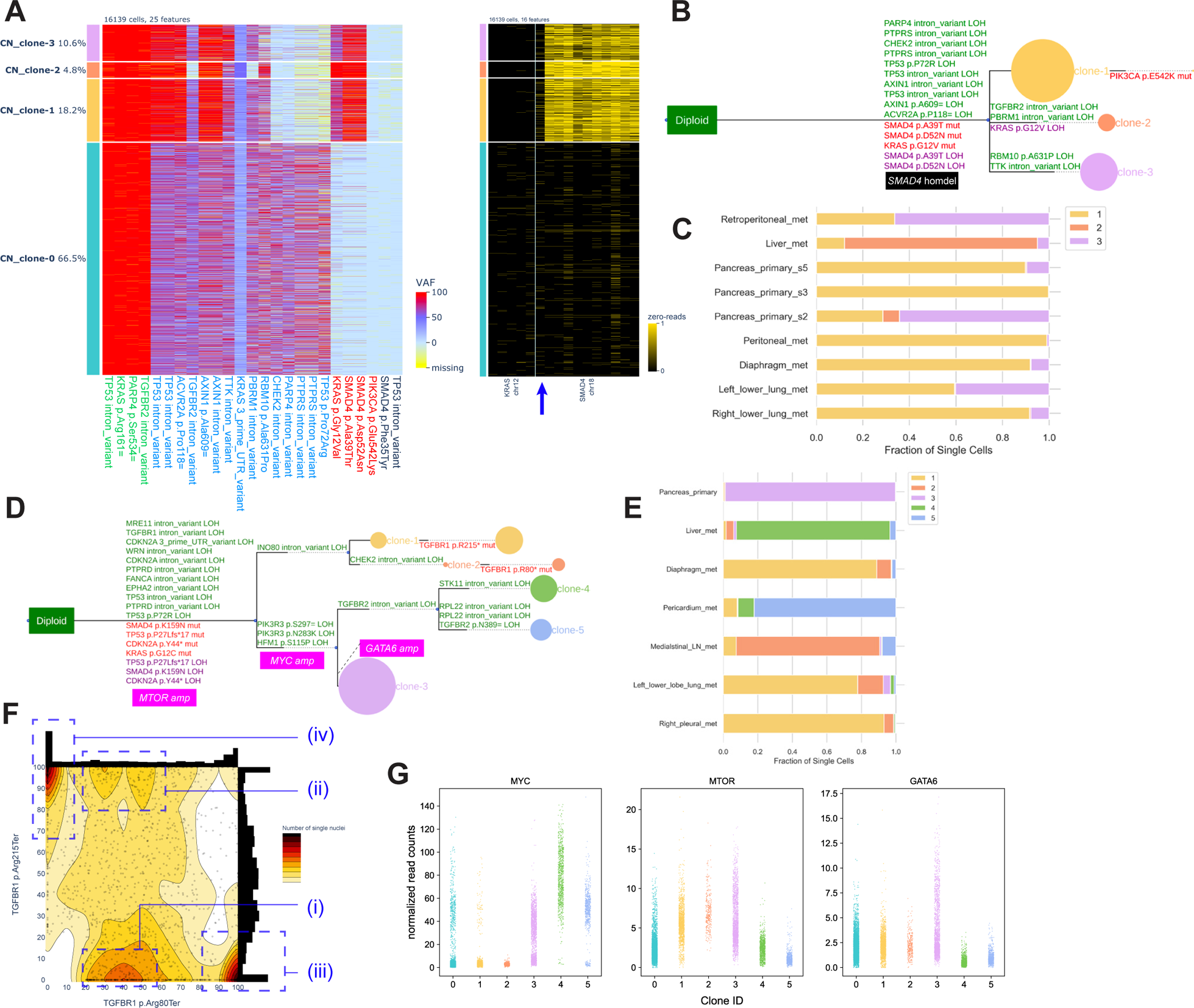
Continuous evolution in pancreatic cancer. **a.** Paired single-cell mutation heatmap and single-cell amplicon heatmap of mutations and genes of interest in PC11. The one SMAD4 amplicon which contained the two somatic mutations is pointed with the arrow. **b.** Phylogeny of PC11. **c.** Clone proportions at each sample site of PC11. **d.** Phylogeny of PC10. **e.** Clone proportions at each sample site of PC10. **f.** Distribution of VAF of the two TGFBR1 mutations of interest across single cells of PC10. The density is labeled as colors and contours, with darker colors indiating higher density of single cells. The four major populations of interest are circled and labeled. **g.** Distribution of normalized read count of MYC, MTOR, GATA6 across single cells of 1 normal clone (clone 0) and 5 tumor clones of PC10.

The continuous evolution also manifests as LOH of somatic driver mutation loci in PC10. Two *TGFBR1* nonsense mutations were subclonally gained in one of the two main lineages of metastatic clones (**Figure 4D)**. The fact that clones of that branch migrated to four spatially distant metastatic sites (**Figure 4E**) indicated that they likely came from a subclone selected for by the primary site rather than any metastatic site. Although fast-ConDoR inferred that the two mutations were gained in different populations (**figure 4D)**, focal analysis of the subclone identified four sub-populations: (i) cells heterozygous for *TGFBR1* p. R80* (mutation “A”) (ii) cells heterozygous for A, homozygous for *TGFBR1* p.R215* (mutation “B”) (iii) cells homozygous for A (iv) cells homozygous for B (**Figure 4F**). Complicated by chromosomal gain/loss, it was difficult to model accurately the evolution path; yet the presence of subclone (ii) indicated that the two mutations were present on different chromosomes in the same set of cells at one time in the tumor’s evolution history, and then the likely scenario was that chromosomal loss resulted in the LOH genotype for either mutations, hence subclones (iii) and (iv).

Aside from deletions, subclonal focal amplification was observed in advanced clones. In the tumor clones of case PC10, the *KRAS* mutant locus had allelic imbalance (single-cell VAF > 0.6); one clone had total loss of the WT allele (**Supplementary Figure 3D**). Coupled with the total copy number gains (**Supplementary Figure 3E**), this suggests that the mutant *KRAS* allele at least tripled in clone 2.

In case PC11, *MYC’*s total copy number (CN) state stayed in the normal range in clones 1 and 2 in the upper branch (**Supplementary Figure 4A)**; for clones 3, 4, 5 in the lower branch, MYC was uniformly amplified to 20+ (**Figure 4G**). Such an amplification pattern was inferred to be caused by extrachromosomal DNA (ecDNA)^23^ (**Supplementary Figure 4B, C**). We interpret the single cells in the normal clone 0 with high *MYC* CN state to be tumor cells that our algorithm could not partition out due to the high total number of cells analyzed (n=8820), because they showed positive signal for the main tumor drivers as well **(Supplementary Figure 4D).** Clones 1 and 2 in the upper branch had *MTOR* amplification instead. Intriguingly, clone 3, exclusively present in the primary tumor sampled, showed amplification signal for both *MYC* and *MTOR,* while also a uniquely high CN state in *GATA6*. Particularly, two populations, with and without high CN state of *MYC*, seem to coexist in clone 3. This suggests that metastatic clonal heterogeneity was well represented at the primary site.

Taken together, through snDNA-seq we observed continuous genomic evolution through copy number loss and/or amplification towards particular genotypes. The driver events acquired and fixed early in tumor development likely culminate in increased genomic instability, which allowed for such CNV events to occur faster than clocklike SNV events^24^ and allowed for rapid adaptation as the tumor invades and metastasizes to foreign environments.

### 4. Evolutionary dynamics through metastasis

Given the knowledge that CNVs are the main driver of ongoing evolution and thus clonal heterogeneity in pancreatic cancer, we next analyzed how this heterogeneity manifests in the context of metastasis. Among 11 patients of MSK’s last wish program (LWP) for research autopsy, we collected in total >= 3 samples per common metastatic site - peritoneum, liver, lung, lymph node (LN), diaphragm - and matched primary samples when available (**Figure 5A**).

**Figure 5:**
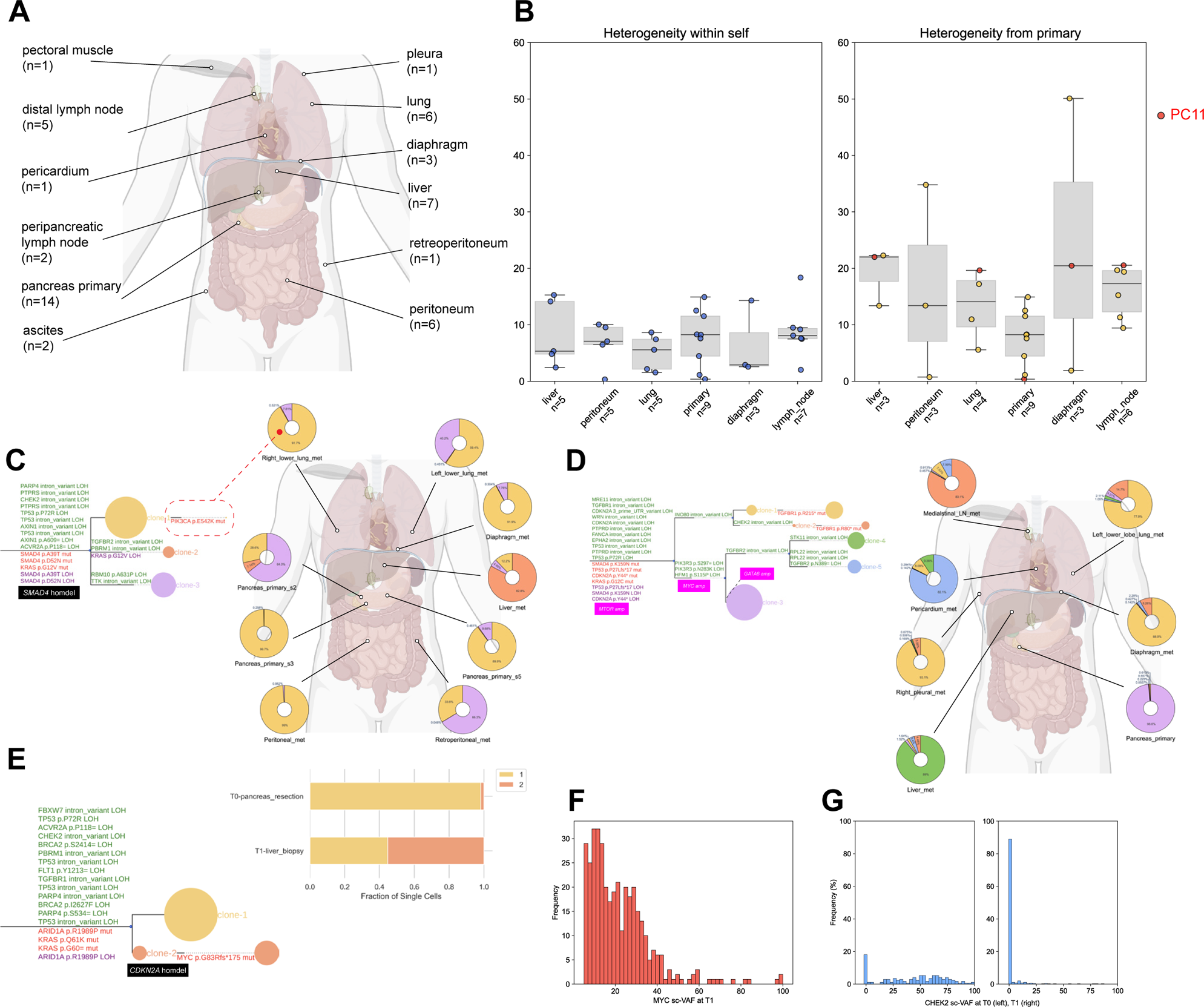
PDAC evolution through metastasis and time. **a.** Distribution of metastatic samples collected by anatomical location. **b.** Distribution of (left) intra-primary/metastatic site clonal heterogeneity and (right) metastatic site vs primary heterogeneity across different anatomical sites. For PC11 (c) and PC10 (d), phylogeny (left) and clonal proportion of each primary/metastatic site, mapped anatomically. **e.** PC20 phylogeny (left) and clone proportion in each sample (right). **f.** Distribution of VAF of PC20’s MYC mutation in cells at the second timepoint (T1). **g.** Distribution of VAF of the germline mutated CHEK2 locus in cells at T0 (left) and T1 (right) of PC20.

We first asked if there is any specific organ selected for mutations to a specific pathway, but did not find any pattern, as bulk studies suggested^25^. Next, we investigated how organ sites correlated with clonal patterns: specifically, if any site would have a stronger selective pressure and sculpt for (1) a more homogeneous clonal composition or (2) clones more advanced compared to the primary site. To answer these two questions, we calculated (1) intra-site heterogeneity and (2) primary-vs-metastasis difference by averaging inter-cell CNV distances between single cell populations from (1) the same site or (2) two separate sites (**Methods**). There was significant inter-patient variability for both metrics (**Figure 5B**). PC10 contributed high primary-metastasis heterogeneity across sites (**Figure 5B**, highlighted in right panel). This is likely due to the peculiar genomic instability which sculpted the complex CNV evolution paths described in the previous section (**Figure 4E**). Two metastases in two other patients also had significant differences from their respective primary site: PC16’s peritoneal metastasis and PC08’s diaphragm metastasis. While the former was mostly caused by preservation of less evolved clones in the primary site (**Supplementary Figure 5A**), the latter was likely caused by elevated chromosomal instability generally, which resulted in the many homdel described previously **(Figure 3D**). These diverse storylines indicate that there is no general rule to organ-specific selective pressure which would make one organ prone to more homogeneous/advanced clonal composition.

Next, we investigated if organ sites proximal to each other tend to have more similar clonal composition, in the three most extensively sampled cases: PC09 (7 sites), PC10 (7 sites), PC11 (9 sites). PC09 and PC11 both presented with comb-like phylogeny indicating high-degree clonal sweep before metastasis and low inter-clone distance. This entailed high spatial homogeneity in these two cases and therefore no correlation between spatial location and clonal composition (**Figure 5C, Supplementary Figure 5B**). An exception might be that PC10’s right lung lower lobe metastasis had a unique subclonal *PIK3CA* p.E542K mutation (**Figure 5C, dashed red circle)**, although orthogonal bulk sequencing found it in both the lung site and another liver metastasis sample not present in this snDNA-seq cohort, suggesting that it was likely pre-metastatic (**Supplementary Figure 5C**). PC11, on the other hand, presented a clearly bifurcating phylogeny with two groups of metastatic sites each carrying clones from only one branch. Upper branch: mediastinal LN, right pleura, left lung lower lobe, diaphragm; lower branch: pericardium and liver (**Figure 5D**). This again demonstrated no correlation between spatial proximity and clonal similarity.

### 5. Evolutionary dynamics through time

Finally, we performed a small pilot study of snDNA-seq as applied to longitudinally collected samples from four patients taken in association with the <u>T</u>racking <u>o</u>f <u>P</u>ancreatic <u>C</u>ancer Regressi<u>o</u>n <u>a</u>nd Resistance (TOPCOAT) program at our institution. Three patients had metastatic disease (stage 4) at diagnosis (T0), and snDNA-seq revealed all relevant drivers in each case. Phylogenies indicated high clonal homogeneity regardless of whether pancreas primary or the liver lesion were biopsied (**Supplementary Figure 6A-C**), consistent with strong selection for the drivers present and one or more clonal sweeps taking place before the first biopsy was taken. For a fourth patient PC20, samples were taken from resection (stage 2A, T0) and a recurrent liver lesion (stage 4, T1). Akin to the other three longitudinal cases, driver alterations including *KRAS* p.Q61Q and *ARID1A* p.R1989P SNVs, *CDKN2A* homdel and many LOH events involving *TP53, TGFBR1* and the germline heterozygous *BRCA2* p.I2627F locus, were fixed at T0 (**Figure 5E).** However, among 450 *KRAS* p.Q61K positive cells at T1, 164 also showed a subclonal *MYC* p.G83Rfs*175 mutation (sample-specific CCF = 0.364), which was completely missing from T0 (**Figure 4E, Supplementary Figure 6D**). The *MYC* mutation had a single-cell VAF (sc-VAF) densely distributed <0.2 (**Figure 5F)**, which, coupled with a total copy number gain at the gene (**Supplementary Figure 6E**), suggests a mutation on the minor allele after total copy number gain. Although the germline CHEK2 SNP locus was inferred by our tree inference algorithm to undergo clonal LOH at T0, we noted that its sc-VAF distribution in tumor cells suggested a subclonal LOH at T0, which dominated at T1 (**Figure 5G)**, which likely corresponds to a focal CNV event after the large-scale CNVs that resulted in the many other LOHs mentioned above.

## Discussion

We presented here a genetic analysis of 24 pancreatic cancers using a new targeted snDNA-seq technique, and a new suite of accompanying computational analysis tools. To our knowledge, this is the first large-scale application of this technique on clinical solid tumor samples. We believe that it supports high utility in the clinical setting for two reasons. First, the wet-lab workflow is largely automated and accommodates even the more challenging clinical samples such as extremely small frozen biopsy samples. One exception is that it cannot perform on formalin-fixed tissues based on testing by both our group and another group, possibly due to DNA damage caused by the fixation process. Second, given single-cell studies’ potential to bring about new biological insights^26^, a targeted workflow enables high-depth sequencing of actionable genes/pathways and yields more clinically relevant information than low-depth pan-genome methods^27,28^, as demonstrated by the success of MSK Integrated Mutation Profiling of Actionable Cancer Targets (IMPACT)^29^. This will become increasingly useful as we are entering the era of precision medicine for pancreatic cancer with the advent of PARP inhibitors for cases with HRD genotype/phenotype and *KRAS* inhibitors for cases carrying oncogenic *KRAS* mutations. Nonetheless, we acknowledge a major limitation of the snDNA-seq method is that, due to its targeted nature, it mostly allows for granular studies of already known targets instead of discovering new targets. Although we curated our targeted panel based on comprehensively curated bulk WES/WGS data of the selected patients, a small fraction of important genetic events was not covered.

The major concepts of pancreatic cancer genetic evolution, such as the main genetic drivers, the genetic progression model, the clonal sweeps before diagnosis and metastasis and the intratumoral heterogeneity driven by CNVs, have been delineated by bulk studies^15,30,31^. Nonetheless, we believe our high-resolution snDNA-seq elucidates the closest-to-ground-truth *baseline* of pancreatic cancer in such contexts as early/late diagnosis, metastasis and non-targeted treatments. While validating the genetic progression model, we refined it with more granular features such as the subclonal occurrence of alteration to *CDKN2A* and *SMAD4,* convergent evolution towards TGF-β pathway inactivation, and continuous evolution through CNVs that are hard to capture by bulk sequencing. Notably, in all six gBRCA2 pancreatic cancers, we discovered varied timing of biallelic *BRCA2* inactivation. It is conceivable that this might explain the heterogeneous treatment response in patients with similar gBRCA2 genotype identified by bulk sequencing^32^, and future studies to confirm or refute this are warranted. As more trial data of targeted treatments start to accumulate, it would be important to compare those against our baseline to identify resistance mechanisms.

## Methods

### Ethics statement

Use of samples used in this study was approved by the institutional review board at Memorial Sloan Kettering Cancer Center (under protocols #06-107, #15-021, #15-149).

### Patient sample collection and preprocessing

Patient samples used in this study consist of three categories-multiregionally sampled surgical resection of primary pancreatic cancer, multiregionally sampled autopsy of metastatic pancreatic cancer and longitudinally sampled biopsies of pancreatic cancer. Each participating patient provided written consent and was not compensated. Detailed patient and sample information is summarized in **Supplementary Table 1**.

For multiregional sampled surgical resections: treatment naïve patients with tumors ≥2 cm on cross-sectional imaging were identified preoperatively. A single cross-sectional piece of tumor was sampled sequentially using a cartesian coordinate system with 0.6 cm × 0.6 cm grid, with 3– 5 samples obtained from each tumor. Adjacent normal pancreas or duodenum was also collected. All samples were stored at −80 °C until use.

For multiregional sampled autopsies: all patients had a premortem diagnosis of PDAC based on pathological review of resected biopsy material and/or radiographic and biomarker studies. Rapid autopsy was performed following our previously described workflow^19^.

The above two types of tissue samples were embedded in optimal cutting temperature (OCT) compound, stained with hematoxylin and eosin (H&E) and reviewed by a gastrointestinal pathologist (S.U.) to ensure total cellularity and tumor purity. Normal samples were reviewed to confirm that no contaminating cancer cells were present.

For longitudinally sampled biopsies, tissue cores were taken by a 22-gauge needle at endoscopic ultrasound (EUS) for primary tumors, or an 18-gauge needle at interventional radiology (IR) for metastatic lesions, and flash frozen until use. The biopsy samples were significantly smaller than the above two sample types, often invisible to naked eyes, precluding them from being pathologically reviewed for cellularity of tumor purity. Tissues stored in collection tubes were suspended in nuclei storage buffer (S2 Genomics) for transferring directly into the Singulator machine (S2 Genomics) for single nuclei extraction.

Additionally, as we were initially concerned that the small volume of biopsies would not be sufficient for snDNA library construction, we expanded one case (PC16)’s 4 of 6 samples into organoids to increase the cell amount. The organoid workflow was as described in Hayashi et al.^33^

Worth noting is that microdissection was omitted so that the resulting snDNA libraries were mostly mixes of tumor and normal cells.

### Bulk sequencing library preparation, sequencing, and bioinformatics

Genomic DNA was extracted from each tissue using the phenol-chloroform extraction protocol or QIAamp DNA Mini Kits (Qiagen). DNA quantification, library preparation, and sequencing were performed in the Integrated Genomics Operation (IGO), where Illumina HiSeq 2000, HiSeq 2500, HiSeq 4000, NovaSeq 6000, NovaSeq X platforms were used to target sequencing coverages of >80× for WGS samples and >150× for WES samples.

Bioinformatics analysis including calling germline/somatic single-nucleotide variant (SNV), copy number variation (CNV), structural variant (SV), estimating mutational signature, scoring HRDetect^34^ were done using the Time-Efficient Mutational Profiling in Oncology (TEMPO), the MSK Center for Molecular Oncology (CMO) Computational Sciences (CCS) research pipeline. Its code repository is available at https://github.com/mskcc/tempo and documentation at https://cmotempo.netlify.app/.

### Single-nucleus DNA sequencing library preparation, sequencing and bioinformatics

#### Nuclei extraction from frozen tissue, counting, QC, cryopreservation

Single nuclei from OCT-embedded snap-frozen primary tissue samples were extracted, counted, quality-controlled and cryopreserved following our published protocol^14^.

#### Panel design for single-cell targeted library preparation

The panel was designed as an expansion of our previous 186-amplicon panel^14^ with the following goals:

1. To unbiasedly cover genes/genomic segments hypothesized to undergo convergent evolution in PDAC, constituting three main categories:

1. canonical PDAC drivers: *KRAS, TP53, SMAD4, CDKN2A*
2. components of the transforming growth factor beta (TGF-β) pathway, including receptors, mediators and downstream effectors: *TGFBR1, TGFBR2, ACVR1B, ACVR2A, BMPR1A, BMPR1B, SMAD2, SMAD3, SMAD7, ARID1A, ARID1B, ARID2*.
3. The *BRCA2* gene, which is of particular interest to PDAC precision medicine.
2. To study genes frequently affected by CNV in PDAC, including *MYC, GATA6, BAP1, MUS81* and *KAT5.* 4+ amplicons were given per gene. At analysis, amplicons for the same gene were constrained to have the same ploidy to obviate the PCR amplification noise of single amplicons and get more confident absolute ploidy calls.
3. To study other recurrent mutations in PDAC, which came from curation of two databases:

a. Bulk whole-exome/-genome sequencing of PDAC autopsy, resection, derived-organoid banks consisting of 100+ patients collected by the Iacobuzio Lab and the MSK David M. Rubenstein Center for Pancreatic Cancer Research.
b. MSK IMPACT sequencing of PDAC attained from MSK cbioportal^35–37^.

The final design was 596 amplicons covering 253 genes’ UCSC canonical exon transcripts. The full list of amplicons and more details are provided in **Supplementary Table 3.**

#### Library preparation and sequencing

Library preparation and sequencing were done as described in our previous paper, except that for this batch of samples we utilized the custom 596-amplicon panel described above.

#### Quality control, cell-calling

FASTQ files for single-nucleus DNA libraries were processed through Mission Bio’s Tapestri pipeline with default parameters. Briefly, it trims adapter sequences, aligns reads to the hg19 genome (UCSC), assigns reads to cell barcodes. The CellFinder module then filtered for barcodes corresponding to “complete cells/nucleus” based on total read completeness (>8 * number of amplicons) and per-amplicon read completeness (>80% data completeness for working amplicons, which are defined as amplicons with > 0.2*mean of all amplicon reads per qualified barcode).

#### Mutation calling

As a first pass, with aligned single-nucleus DNA reads (binary alignment maps, BAMs), Mutect2 (GATK 4.2.5.0) was used for mutation calling and filtering in each single-cell BAM. Preliminary technical filters were applied, including: *base_qual/low_allele_frac/weak_evidence/slippage/multiallelic/clustered_events*. Required minimum depth was 4 reads and variant allele frequency (VAF) was 0.2. Still, we observed many called mutations present in small numbers of single barcodes, which are likely false positives from PCR/sequencing errors^38^. We concatenated mutations called in each single cell, and with the belief that the false positive rate of a mutation decreases exponentially with the number of barcodes it is detected in, we only considered mutations present in >= 3 single cells.

As a second pass, we used the mutation list from the first pass for genotyping in each single-cell BAM with *bcftools mpileup,* outputting a mutation by single-cell matrix that has information for each mutation in each single cell.

Further, to enable *de novo* mutation calling, we took the following steps of quality control, inspired by bulk DNA sequencing bioinformatics.

#### Detecting rare variants leveraging multiregional sampling

Sampling multiple regions of a tumor enables mutations that have a large clonal fraction in one sample but lower than detection threshold fraction in another to be captured in the latter.

Therefore, mutations from all samples of the same tumor were pooled and genotyped in all single nuclei with *bcftools mpileup*.

#### Filtering by relative prevalence and mutational signature

In both neoplastic and normal samples taken from different organ sites, we noticed a significantly similar mutational signature, signified by a large proportion of T>C variants, enriched in mutations <0.5% single-cell prevalence detected by this snDNA-seq technology (**Supplementary Figure 1B**). As different types of human tissues have been shown to display distinct mutational signatures and the signature did not seem to match any of themcos^39^, we deemed it likely artifactual and decided to filter out all mutations below 0.5% single-cell prevalence. T>C variants in the 0.5% to 1% were also filtered.

#### Assembly of a panel of normal

A panel of normal (PON), proven effective in eliminating library artifacts in bulk sequencing, was assembled for snDNA-seq from three sources:

1. 8 unrelated normal pancreas samples were subject to the same snDNA-seq sequencing and mutational calling process (until filtering by prevalence). Variants called in >= 4 unmatched normal samples were discarded.
2. Variants present in a bulk WES PON (gs://gatk-best-practices/somatic-b37/Mutect2-exome-panel.vcf) were discarded.
3. SnDNA-seq SNV lists of all tumor samples studied here were concatenated, and variants called in >50% samples, except for those in known hotspot (*KRAS* codon 12), were discarded. Manual curation proved this effective in removing low-prevalence SNVs likely caused by library error, and confirmed that no driver mutations identified by matched bulk sequencing were discarded.

To retain real SNPs/SNVs with high prevalence in the population, variants with a frequency > 0.01 in the 1000 Genome Project were whitelisted.

#### Additional filters for phylogenetic analysis

As our phylogenetic analysis relies on high-density single-cell VAF signal of mutations, those with >0.3 biallelic dropout (ADO) rate in any sample were discarded.

#### Estimation of the highest resolution of bulk sequencing and Tapestri in calling somatic mutations

We referred to Frankell et al.^13^ for a state-of-the-art bulk sequencing bioinformatics pipeline. As described in their method: *“A SNV was considered a true positive if the variant allele frequency (VAF) was greater than 2% and the mutation was called by both VarScan2, with a somatic P ≤ 0.01, and MuTect. Alternatively, a frequency of 5% was required if only called in VarScan2, again with a somatic P ≤ 0.01. Additionally, the sequencing depth in each region was required to be ≥30, and ≥10 sequence reads had to support the variant call.”* From here, we took the most relaxed threshold 2% VAF, and assumed that the driver mutations of interest have loss of heterozygosity in cancer cells. We arrived at 2% single-cell prevalence as bulk sequencing’s highest resolution of somatic mutation calling.

For Tapestri, as mentioned in a previous section, because it can to pick up any mutation present in any single barcode, we estimated its sensitivity to be 1 / 1858 (mean per-sample library size) ≈ 0.05% single-cell prevalence. However, due to the noise profile we observed below 0.5%, we estimated its specificity to be limited to that number.

### Single-nucleus DNA sequencing genetics analysis

#### Calculating cancer cell fractions

With snDNA-seq, because we can observe VAF of each SNV in each single cell, we could calculate CCF directly by designating one clonal population as cancer, and measure the number of cells within that population that shows a confident signal for each mutation.

We first genotyped each mutation in each cell based on hard thresholding: a mutation is considered positive in a cell given >= 3 mutant reads, >= 0.2 VAF and >= 8 total depth.

To define the cancer clonal population, we assumed *KRAS* as the earliest driver event^40^ and considered those positive for *KRAS* as cancer. Thus, *KRAS* mutations would have CCF = 1 for all cases that carry them, and another mutation *i* would have *CCF* = *N_i_* / *N_KRAS_*, where N is the number of cells carrying each mutation. For those cases that did not have a *KRAS* mutation, we designated the union of all non-diploid clones, as inferred by fast-ConDoR as cancer.

Amplifications and homozygous deletions’ CCFs were calculated as the proportion of their corresponding clones as inferred by Tapestri-CN. LOHs’ CCFs were calculated as the proportion of their corresponding clones as inferred by fast-ConDoR.

#### dN/dS

We used the trinucleotide context-dependent substitution model as implemented by *dndscv*^41^. Due to the limited size of our panel and cohort, we did not implement the variational mutation rates across different genes and tumors. For each gene of interest, its dN value would be calculated as the number of missense/nonsense mutations normalized by the total number of possible missense/nonsense mutations on our panel; dS would be a similar calculation albeit for synonymous mutations.

### Single-nucleus DNA sequencing phylogenetics analysis

#### Copy number calling from single-cell DNA-sequencing data

We developed a new tool, “Tapestri-CN” to infer the copy number profiles of individual cells from the single-cell DNA-sequencing data. The tumor sample from each patient is composed of multiple cancer clones with distinct copy number profiles in varying proportions. As such, we model each tumor sample as a mixture of multiple copy number profiles with unknown proportions, where the read count data is given by a Negative Binomial model. We derive an expectation maximization algorithm to infer the copy number clones, their proportions and the assignment of each cell to one of the copy number clones in each tumor sample. Details of the model and testing/validation process are provided in **Supplementary Methods**. The code repository is available at https://github.com/haochenz96/tap_cn_calling.

#### Single-cell clone phylogeny inference and refinement

We modified our previously published method ConDoR^18^ to make it scalable to our dataset, which had cases with 10,000+ single cells as all samples from the same patient were merged. The modifications to achieve scalability can be found in **Supplementary Methods**. The updated method Fast-ConDoR takes the following as input: (i) variant and total read counts of mutation (SNPs or SNVs) in each cell, (ii) copy number clustering of cells into distinct copy number clones, and (iii) annotation of the mutations into two groups – germline SNPs and somatic SNVs. The annotation of mutations is required because while SNVs are somatic mutations that may be gained and lost during cancer evolution, SNPs are germline mutations that are only impacted by LOH events.

We obtain the annotation of the mutation measured from snDNA-seq data by comparing them with mutations derived from bulk sequencing data of matched normal and tumor samples. Since homozygous SNPs are not informative for cancer phylogeny inference, mutations with pseudo-bulk VAF > 0.9 in snDNA-seq were filtered out. Upon testing fast-ConDoR, we found a set of loci recurrently called as LOH in non-neoplastic clones (i.e. cells without driver mutations such as *KRAS, TP53*) in multiple patients, likely arising due to artifactual allelic imbalance at corresponding amplicons caused by PCR. These loci were manually selected and filtered out.

Fast-ConDoR produces a tree where the leaves represent the sequenced cells and the edges indicate gain or less of SNVs and SNPs. To avoid overfitting the data, we merge copy number clones with low proportions (i.e. comprise less than 0.1% of total cells) and do not have any SNVs. This was implemented because based on observation of our samples, such clones always had only likely false positive LOHs called, thus could not be confirmed. Branch lengths are calculated as described in the next section.

#### Defining driver, finding the trunk of the tree

Because we designed our panel to target only genes critical pancreatic cancer development and progression, we considered all functional mutations (non-synonymous, intron etc.) detected by this panel to be driver events.

Based on the pancreatic cancer genetic progression model and our observation of the inferred phylogenies that none had two early-diverging tumor branches, we simply define the trunk to be the first edge that has somatic SNV event when traversing the tree in preorder. For PC22 and PC24 which do not have somatic SNV covered by our panel, their trunk could be trivially found by hand.

#### Measuring tumor heterogeneity at the sample level

For comparing multiregional samples, we sought to quantify tumor heterogeneity at the sample or site level. We define the spatial heterogeneity metric within the same site or across different sites of the same patient as the average phylogenetic distance between two randomly drawn cells from the same spatial site or two different spatial sites, respectively. In the following, we describe (i) the phylogenetic distance between the cells in a given phylogeny and (ii) the derivation of heterogeneity of a sample.

Phylogenetic distance between two cells, i.e. leaves of a phylogeny, is the sum of branch lengths along the unique path that connects them. We use the maximum parsimony principle to assign branch lengths to the phylogenies inferred by fast-ConDoR for each sample. Specifically, we assign copy number profiles and SNVs to each internal node of the phylogeny using the Sankoff algorithm^42^. The branch length *l_e_* of each edge *e* in the phylogeny is calculated by combining the difference between the copy number and SNV states of the source vertex *u* and target vertex *v* of the edge as follows,

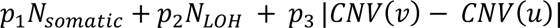

where *CNV*(*v*) is a vector describing the copy number of the internal vertex *v*, *N*_45678’9_ is the number of somatic SNVs gained along the edge and *N*_<=>_ is the number of LOHs that occurred on the edge. *p*_1_, *p*_2_ and *p*_3_ are normalization factors which we set to 100/3, 100/3 and 0.5, respectively.

The average phylogenetic distance between two randomly drawn cells in a sample is given by

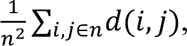

where *d*(*i*, *j*) is the distance between cell *i* and cell *j*.

We quantify the heterogeneity between cells from two samples, say sample A and B, in the same patient as the average phylogenetic distance between a randomly drawn cell from A and another from B. Specifically, the heterogeneity between samples A and B is

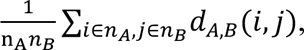

where, *d_A_*_,*B*_(*i*, *j*) is the distance between cell *i* from tumor A and cell *j* from tumor B.

In the same patient, multiple samples of the same general site were combined (e.g. mesenteric nodule and omental nodule would be combined into peritoneal metastasis; multiple nodules at the liver would be combined), so that each data point would represent one site or primary-metastasis pair.

## Data availability

Raw FASTQ files and processed data required to replicate this work will be deposited to European Genomephenome Archive (EGA) shortly after this preprint publication.

## Code availability

The custom bioinformatics pipeline for Tapestri variant calling is available at https://github.com/haochenz96/TapVarCallSmk. The fast-ConDoR pipeline for phylogenetic analysis is available at https://github.com/Bolladeen/full-ConDoR. The downstream analyses to reproduce figures in the manuscript are available at https://github.com/haochenz96/Tapestri_main_manuscript_analysis.

## Acknowledgements

We thank the Integrated Genomics Operations (IGO), bioinformatics core, Center for Molecular Oncology (CMO) of Memorial Sloan Kettering Cancer Center (MSKCC) for their technical support. We thank the Iacobuzio Lab, David M. Rubenstein Center for Pancreatic Cancer Research of MSKCC for their work on sample collection and processing. We thank the Raphael Lab (Princeton University) for their contribution to computational analysis. We thank Alvin Makohon-Moore (Hackensack Meridian Health), Sohrab Shah (MSKCC), Richard White (University of Oxford), Cyrille Delley (University of California, San Francisco), Yichen Wang (Wellcome Sanger Institute), Marcin Imielinski (MSKCC), Marc Arribas-Layton (Mission Bio), Ronan Chaligné (MSKCC), Ignas Masilionis (MSKCC) for their input to the project.

---

**Supplementary Figure 1:**
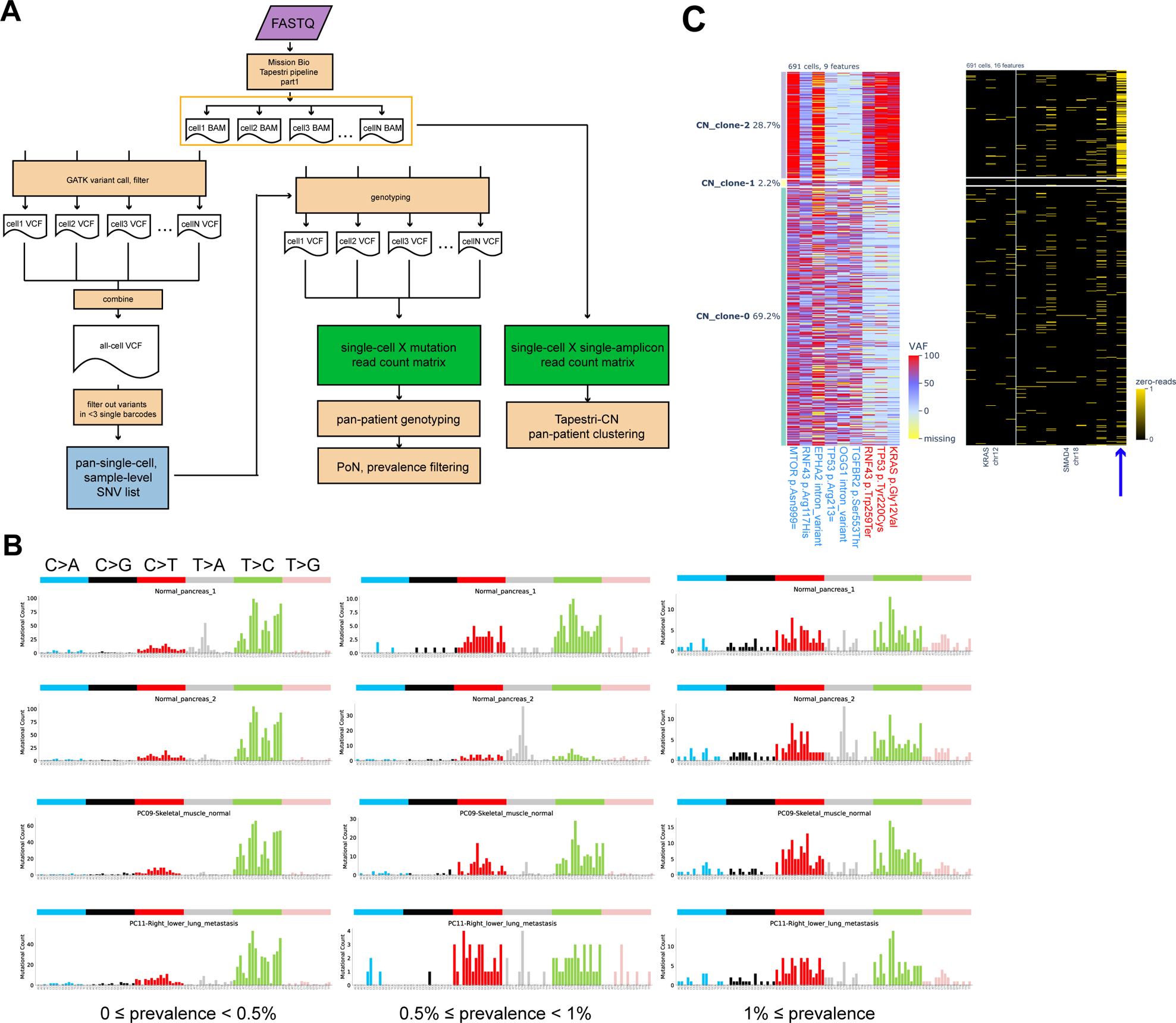
single-nucleus DNA sequencing workflow and output technical details. **a.** Overview of the bioinformatics pipeline, which processes FASTQ files into (1) single-cell by variant matrix (2) single-cell by single-amplicon read count matrix. These two outputs are used for downstream computational analysis to resolve copy number variation (CNV) clones and single-cell clonal phylogeny. **b.** Mutation spectra at varying single-cell prevalence (ordered horizontally) for 4 select samples, including 2 normal pancreatic tissues, 1 skeletal muscle normal tissue and 1 pancreatic cancer metastasis to right lower lung (ordered vertically). Y-axis represents the number of mutations; x-axis represents trinucleotide contexts. Note the enrichment of T>C mutations in lower prevalence mutations across all samples. **c.** For PC18, paired single-cell mutation heatmap, where the heat represents variant allele frequency (VAF) of each mutation in each single cell (left), with single-cell amplicon heatmap, where the heat represents binary indicator of whether an amplicon has zero read or not in each single cell (right). The single cells (rows) are aligned between the two subplots. The one SMAD4 amplicon that likely underwent homozygous deletion (homdel) is pointed by the arrow.

**Supplementary Figure 2:**
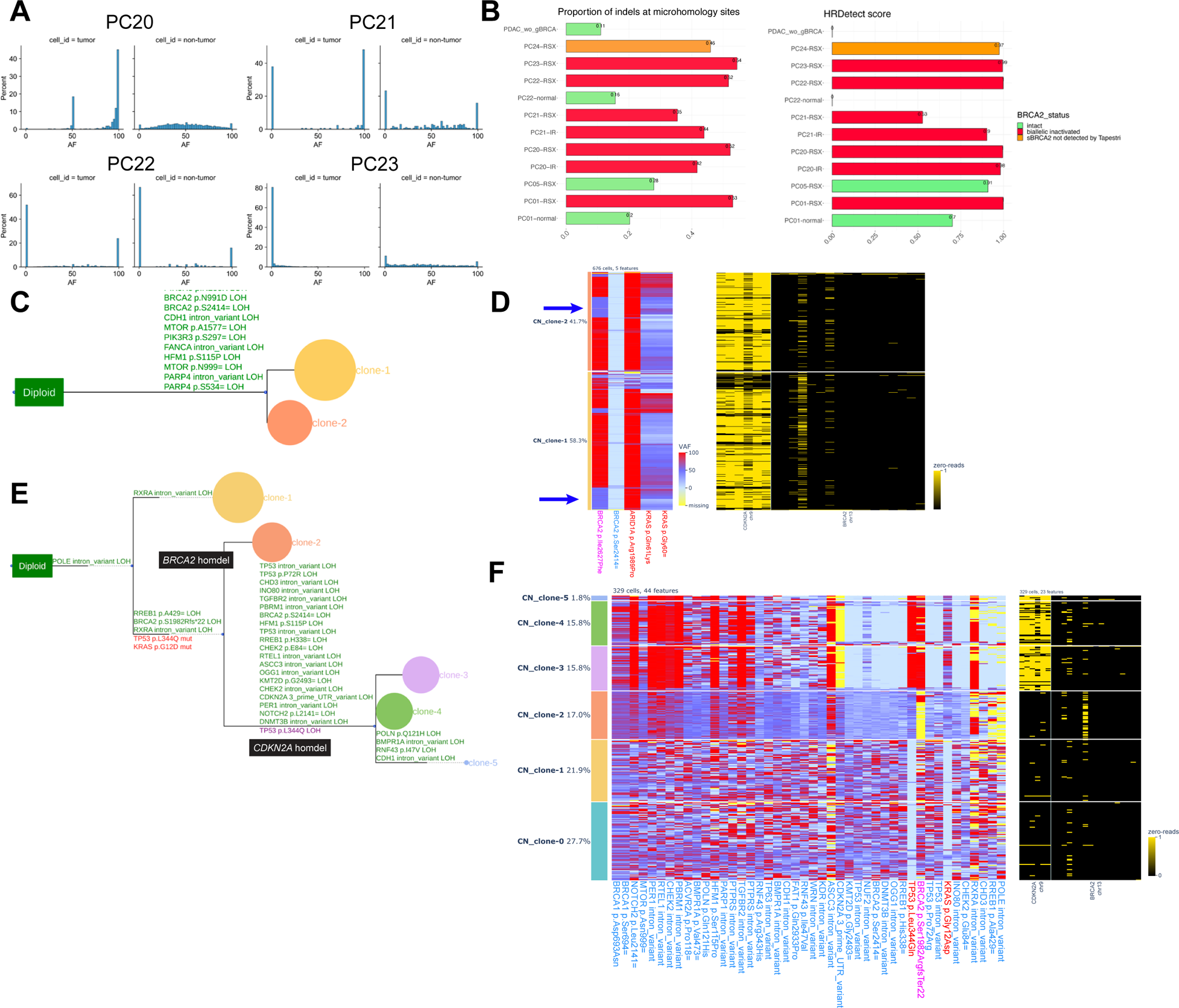
Genomic evolution of pancreatic cancers with germline BRCA2 mutations (gBRCA2) **a.** Variant allele frequency (VAF) distribution of the germline mutated BRCA2 (gBRCA2) locus in single cells of cases PC20, PC21, PC22, PC23. Within each panel, the tumor cell population is on the left and the non-tumor population is on the right. **b.** Homologous recombination deficiency (HRD) characterization of cases with and without biallelic inactivation of BRCA2. Left: proportion of insertions and deletions (indels) at microhomology sites for each case. Right: HRDetect (Methods) score for each case. **c.** Phylogeny of PC23. **d.** For PC20, paired single-cell mutation heatmap and single-cell amplicon heatmap of mutations and genes of interest. Cell populations of interest are pointed with arrows. **e.** Phylogeny of PC21. **f.** Paired single-cell mutation heatmap and single-cell amplicon heatmap of mutations and genes of interest in PC21. The clone of interest is clone 2, where the gBRCA2 locus likely had homdel.

**Supplementary Figure 3:**
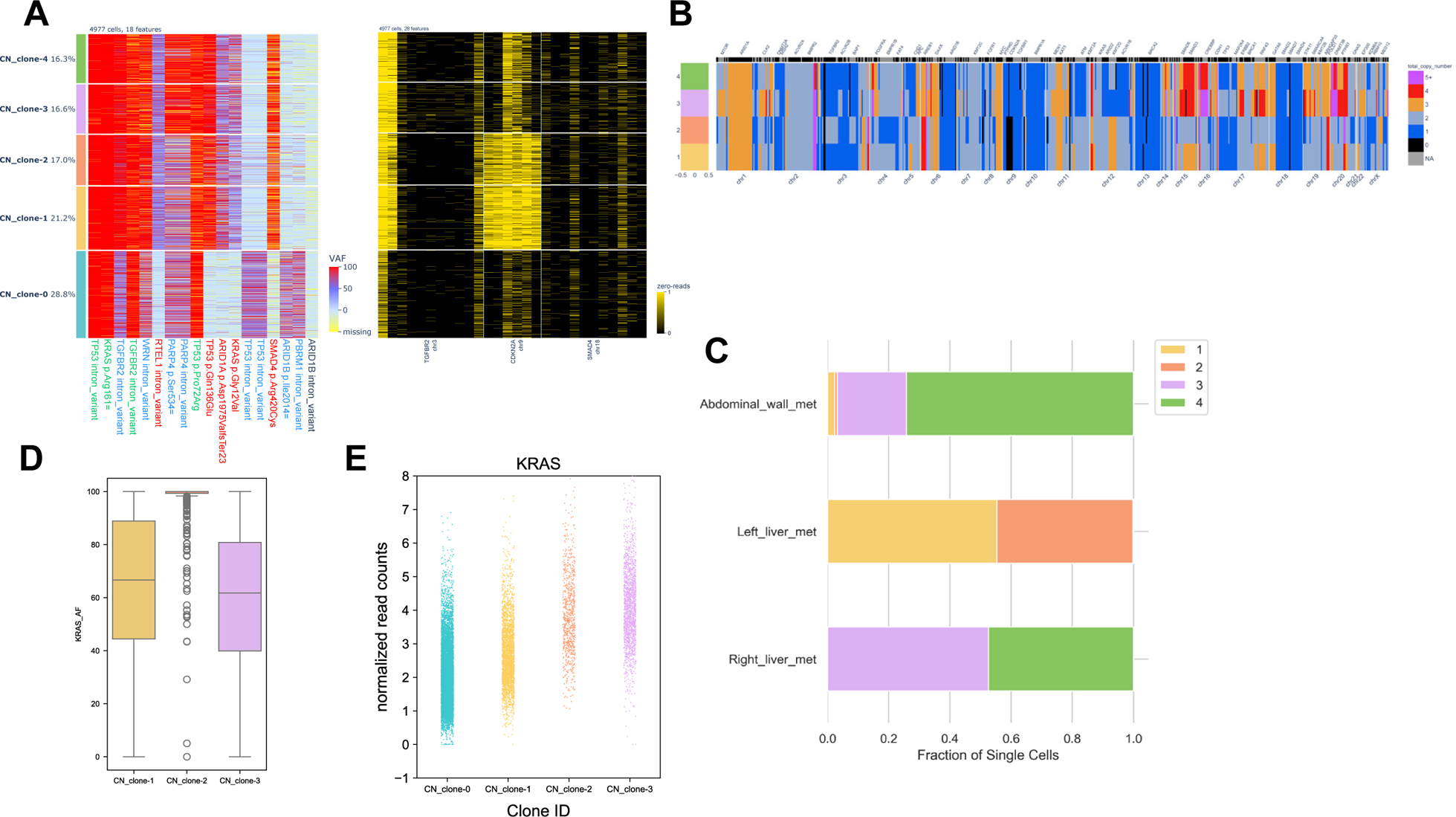
Continuous evolution in pancreatic cancer. **a.** Paired single-cell mutation heatmap and single-cell amplicon heatmap of mutations and genes of interest in PC06. We noted deletions of all 6 CDKN2A amplicons, but were not confident to call subclonal homdel in amplicons #3, 4, 5 because they had deletion signal in the normal tissue of not only this case but multiple other cases, which could likely attribute to primer failure due to high GC content in the region. **b.** Clone total copy number profiles of PC06. **c.** Clone proportions at each sample site of PC06. **d.** VAF distribution of the driver KRAS mutation across cells of 3 tumor clones of PC11. **e.** Normalized read count (Methods) distribution of KRAS in the normal clone (0) and 3 tumor clones of PC10.

**Supplementary Figure 4:**
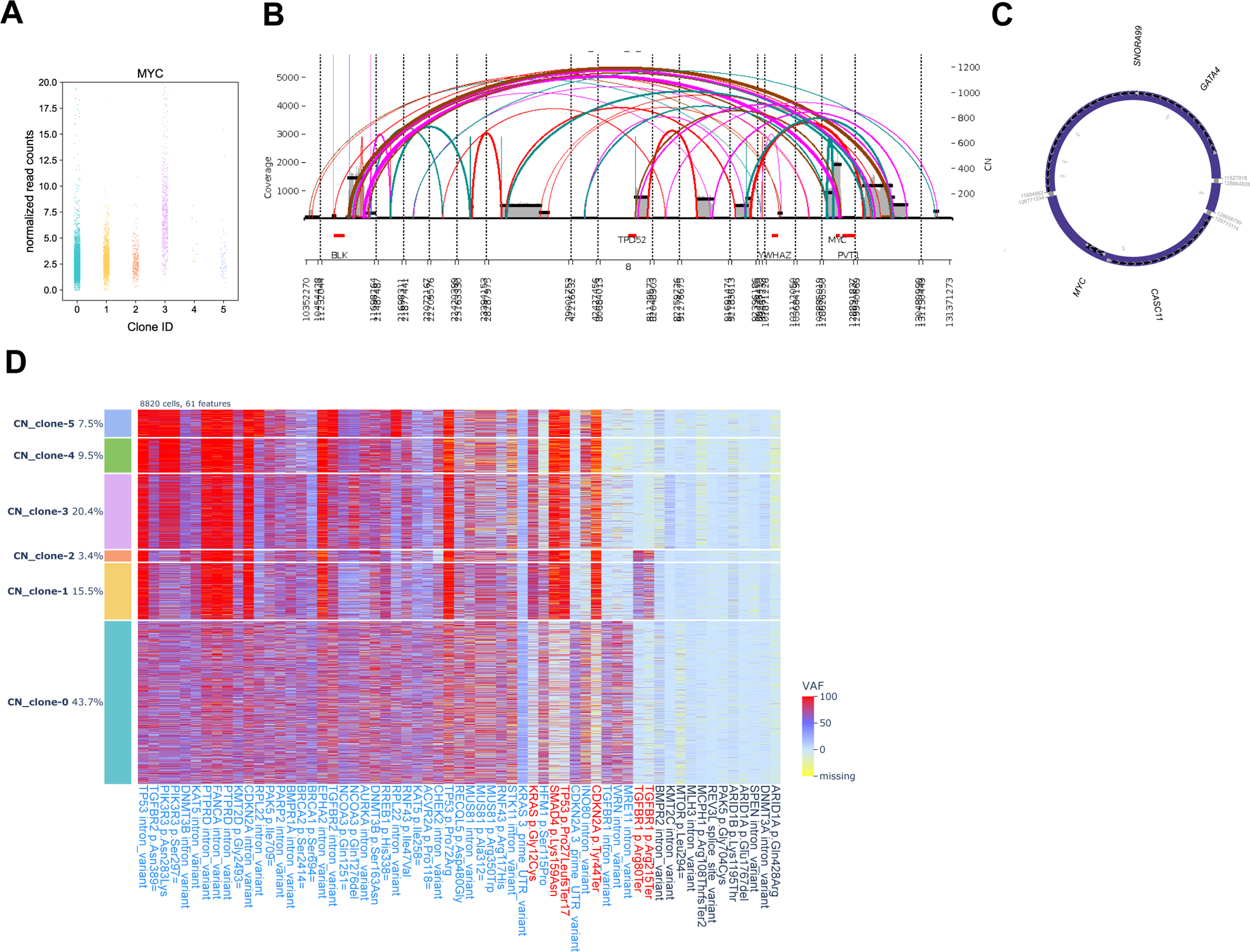
Detailed analysis of PC10. **a.** Distribution of normalized read count of MYC across single cells of 1 normal clone and 5 tumor clones of PC10, with y-axis adjusted to show the lower read count states. **b.** c. AmpliconArchitect (Methods) results of PC10 liver metastasis. **d.** Single-cell mutation heatmap of mutations of interest of PC10.

**Supplementary Figure 5:**
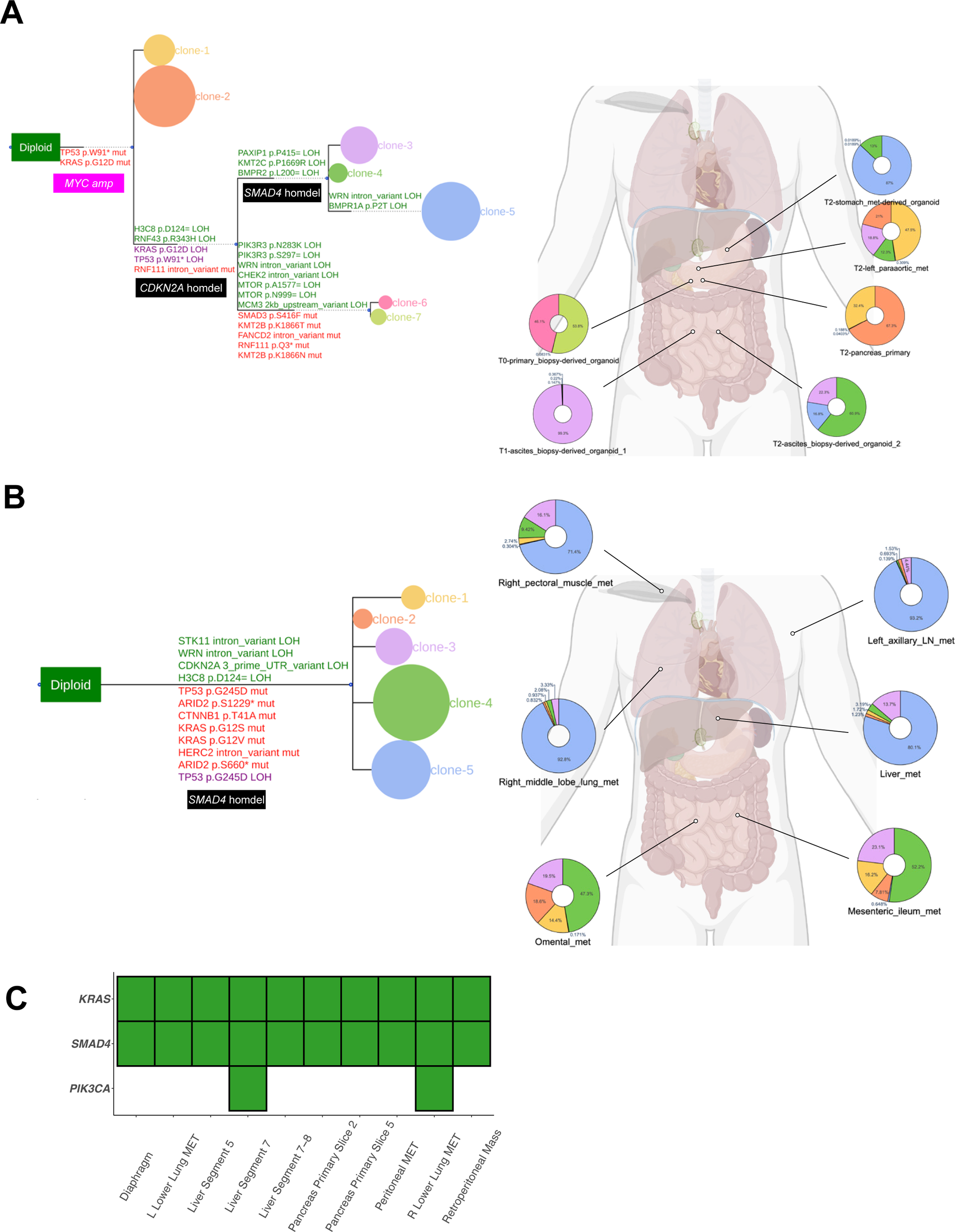
Evolution trajectories of PC09, PC11, PC16 metastases. For PC16 (a) and PC09 (b), phylogeny (left) and clonal proportion of each primary/metastatic site, mapped anatomically. **c.** Based on bulk sequencing, mutation status of the three mutations of interest of PC11 across metastatic sites.

**Supplementary Figure 6:**
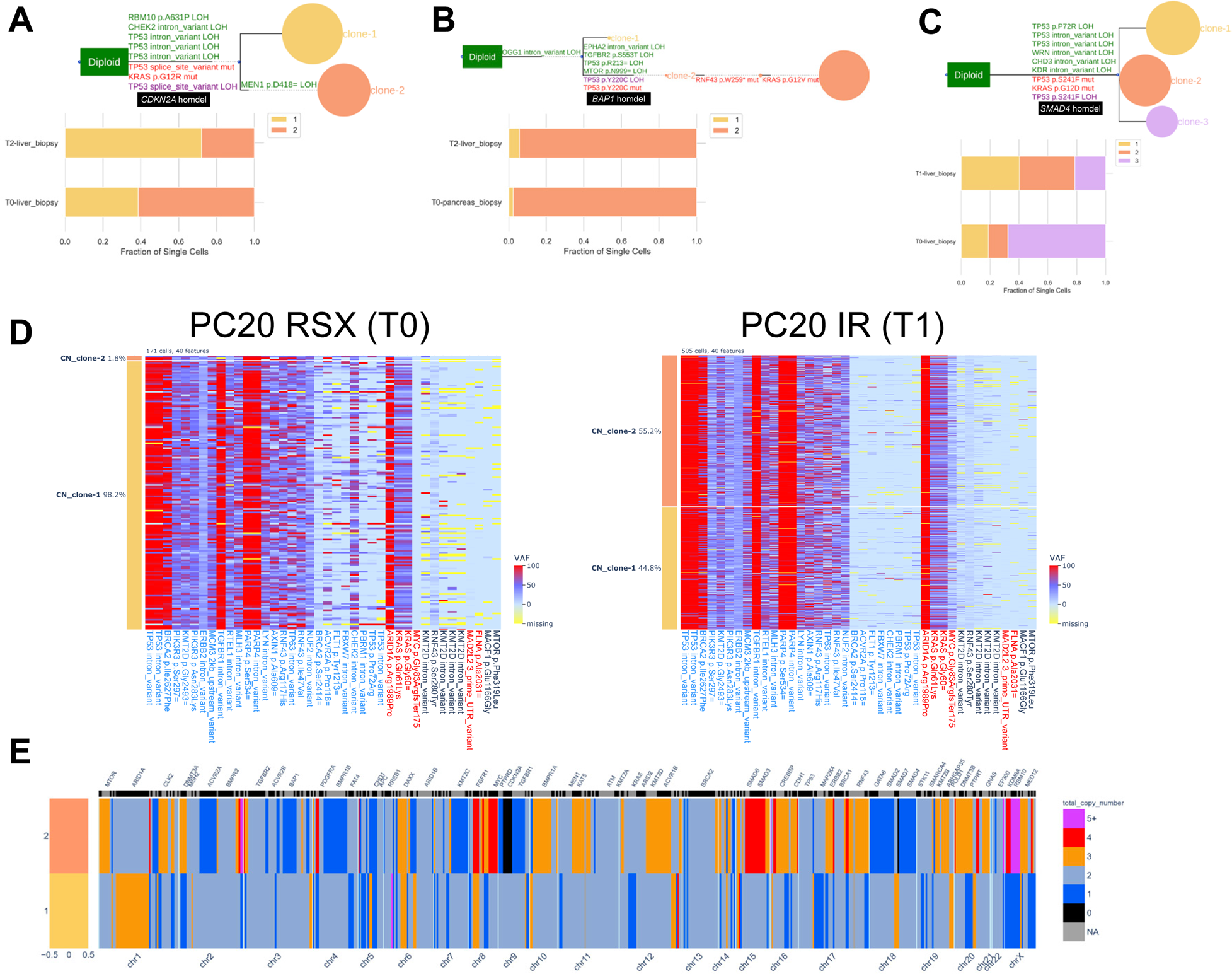
Evolution trajectories of longitudinal samples of PC17, PC18, PC19, PC20. Phylogenies (top) and clone proportions in each sample (bottom) of PC17 (a), PC18 (b), PC19 (c). **d.** Single-cell mutation heatmaps of mutations of interest for samples at two timepoints (T0, left; T1, right) of PC20. **e.** Clone total copy number profiles of PC20.

