## Supplementary Methods for "Genomic evolution of pancreatic cancer at single-cell resolution"

#### 1 Copy number calling from targeted scDNA-seq data

Suppose we measure the number of reads aligned to $m$ genes, each covered by $d$ amplicons, from $n$ cells. We represent this measurement by a read count tensor $R$, where entry $r_{i,j,\ell}$ is the number of reads aligned to amplicon $\ell$ of gene $j$ in cell $i$. Suppose the cancer tumor is comprised of $p$ copy number (CN) clones. We assume that all loci of a gene have the same copy number, except in regions of focal homozygous deletions (homdel), i.e. when the copy number is 0. The copy number clones can thus be represented by two matrices, (1) *copy number matrix* $C$, where entry $c_{k,j}$ is the copy number of gene $j$ in clone $k$ and (2) *homdel matrix* $H$, where entry $h_{k,j,\ell}$ is 0 if amplicon $\ell$ of gene $j$ lies in a region that has been homozygously deleted in clone $k$.

We encode the membership of a cell $i$ to one of the $p$ copy number clones by random variables $Z$, where $z_{i,k} = 1$ if and only if cell $i$ belongs to copy number clones $k$. Note that since each cell belongs to exactly one copy number clone, we have $\sum_{k=1}^{p} z_{i,k} = 1$.

We model the number of reads aligned to an amplicon by a negative binomial model, where the mean of the distribution is proportional to the copy number of the genomic region spanned by the amplicon. Specifically, we model the read count $r_{i,j,\ell}$ for cell $i$ belonging to copy number clone $k$ as follows,

$$r_{i,j,\ell} \mid z_{i,k} = 1 \sim \mathrm{NB}(\mu = h_{k,j,\ell} c_{k,j} f_{j,\ell} T_i, \phi = 1/\beta_{j,\ell}),$$

where $T_i$ is the total number of reads in cell $i$, $f_{j,\ell}$ is the amplicon-specific factor amplification factor of amplicon $\ell$ in gene $j$ and $\beta_{j,\ell}$ is the heterogeneity (dispersion) parameter of amplicon $\ell$ in gene $j$. The amplicon-specific parameters, i.e. $f_\ell$ and $\beta_\ell$ are learned by fitting the above model on cells collected from unmatched normal samples, which were assumed to be diploid, i.e. samples where copy number $c_{i,j} = 2$ for each gene in each cell.

We assume that the cells are sequenced independently. Let $\boldsymbol{\alpha} = (\alpha_1, \ldots, \alpha_p)$ be the mixing proportions of the $p$ copy number clones such that $\sum_{k=1}^{p} \alpha_k = 1$. Let $\boldsymbol{\theta} = (\boldsymbol{\alpha}, C, H)$. The likelihood of observing read count matrix $R$ for a given copy number matrix $C$, homdel matrix $H$ and clone proportions $\boldsymbol{\alpha}$ is as follows,

$$\Pr(R \mid \boldsymbol{\theta}) = \prod_{i=1}^{n} \prod_{j=1}^{m} \prod_{\ell=1}^{d} \Pr(r_{i,j,\ell} \mid \boldsymbol{\theta}),$$

where we assume that the read count measurement of each amplicon in each cell is independent.

We define the copy number calling problem as follows.

**Problem 1.1** (Copy number calling). *For a given read count tensor $R$ for $n$ cells and $m$ genes with $d$ amplicons, find the copy number matrix $C$, homdel matrix $H$ and copy number clone proportions $\boldsymbol{\alpha}$ that maximize the likelihood* $\Pr(R \mid C, H, \boldsymbol{\alpha})$.

We solve the above problem using an expectation maximization algorithm. Specifically, we want to maximize

$$
\begin{aligned}
\Pr(R \mid C, H, \boldsymbol{\alpha}) &= \prod_{i=1}^{n} \Pr(R_i = r_i \mid \boldsymbol{\theta}) \\
&= \prod_{i=1}^{n} \sum_{z_i} P(R_i = r_i \mid Z_i, \boldsymbol{\theta}) P(Z_i) \\
&= \prod_{i=1}^{n} \sum_{k=1}^{p} P(R_i = r_i \mid Z_i = e_k, \boldsymbol{\theta}) P(Z_i = e_k) \\
&= \prod_{i=1}^{n} \sum_{k=1}^{p} \alpha_k \prod_{j=1}^{m} \prod_{\ell=1}^{d} \mathrm{NB}(r_{i,j,\ell} \mid \mu = \rho_{k,\ell} c_{k,j} f_\ell R_i, \phi = 1/\beta_\ell).
\end{aligned}
$$

For brevity, for the remainder of the manuscript, we define

$$
\begin{aligned}
\mu_{i,j,\ell,k} &= \rho_{k,\ell} c_{k,j} f_\ell R_i, \\
\phi_\ell &= 1/\beta_\ell.
\end{aligned}
$$

**EM derivation:** We start by expressing the conditional probability of $Z$ and joint probability of $(X, Z)$, which will be useful later.

$$
\begin{aligned}
P(R, Z \mid \boldsymbol{\theta}) &= \prod_{i=1}^{n} P(R_i \mid Z_i, \boldsymbol{\theta}) P(Z_i) \\
&= \prod_{i=1}^{n} \prod_{k=1}^{p} \left(\alpha_k \mathrm{NB}(r_i \mid \mu_k, \phi_k)\right)^{z_{ik}}.
\end{aligned}
$$

$$
\begin{aligned}
P(Z_i = e_k \mid R_i) &= \frac{P(R_i \mid Z_i = e_k) P(Z_i = e_k)}{P(R_i)} \\
&= \frac{\alpha_k \mathrm{NB}(r_i \mid \mu_k, \phi_k)}{\sum_{\ell=1}^{p} \alpha_\ell \mathrm{NB}(r_i \mid \mu_\ell, \phi_\ell)} \\
&= \frac{\alpha_k \prod_{j=1}^{m} \mathrm{NB}(r_{i,j} \mid \mu_k, \phi_k)}{\sum_{\ell=1}^{p} \alpha_\ell \prod_{j=1}^{m} \mathrm{NB}(r_{i,j} \mid \mu_\ell, \phi_\ell)}.
\end{aligned}
$$

The EM algorithm involves maximizing $Q(\theta, \theta^0)$ in the M-step of each iteration.

$$
Q(\theta, \theta^0) = E_{\theta^0}[\log(P(R, Z \mid \theta)) \mid X].
$$

First note that,

$$
\log(P(R, Z \mid \theta) = \sum_{i=1}^{n} \sum_{k=1}^{p} Z_{ik} \left(\log(\alpha_k) + \log(\mathrm{NB}(r_i \mid \mu_k, \phi_k))\right).
$$

Taking the required expectation, we get the following.

$$E_{\theta^0}[\log(P(X, Z \mid \theta)) \mid R] = \sum_{i=1}^{n}\sum_{k=1}^{p} E_{\theta^0}[Z_{ik} \mid R_i] \left(\log(\alpha_k) + \log(\text{NB}(r_i \mid \mu_k, \phi_k))\right),$$

where

$$E_{\theta^0}[Z_{ik} \mid R_i] = \frac{\alpha_k^0 \prod_{j=1}^{m} \text{NB}(r_{i,j} \mid \mu_k^0, \phi_k^0)}{\sum_{\ell=1}^{p} \alpha_\ell^0 \prod_{j=1}^{m} \text{NB}(r_{i,j} \mid \mu_\ell^0, \phi_\ell^0)} = \gamma_{i,k}.$$

We construct the following Lagrangian

$$\mathcal{L}(\theta, \theta^0) = \sum_{i=1}^{n}\sum_{k=1}^{p}\left(\gamma_{i,k}(\log(\alpha_k) + \sum_{j=1}^{m}\log(\text{NB}(r_{i,j} \mid \mu_k, \phi_k)))\right) - \lambda\left(\sum_{k=1}^{p}\alpha_k - 1\right),$$

where $\lambda$ is a Lagrange multiplier. Setting derivative with respect to $\alpha_k$ to zero together with the constraint $\sum_{k=1}^{p}\alpha_k = 1$ we get,

$$\begin{aligned}\alpha_k &= \frac{\sum_{i=1}^{n}\gamma_{i,k}}{\sum_{k=1}^{m}\sum_{i=1}^{n}\gamma_{i,k}} \\ &= \frac{\sum_{i=1}^{n}\gamma_{i,k}}{n}\end{aligned}$$

We only need to numerically solve for $(\mu_1, \ldots, \mu_k, \phi_1, \ldots, \phi_k)$ by maximizing

$$Q'(\theta, \theta^0) = \sum_{i=1}^{n}\sum_{k=1}^{p}\gamma_{i,k}\sum_{j=1}^{m}\log(\text{NB}(r_{i,j} \mid \mu_k, \phi_k)).$$

We solve this numerically at each iteration of the EM algorithm by brute-forcing all possible values of $c_{k,j} \in \{1, \ldots, c_{\text{max}}\}$. We set $c_{\text{max}} = 20$ for this study. We run the EM algorithm with number $p$ of clones ranging from 2 to 10, and found that usually as $p$ got higher, the number of clones post-refinement (see **Single-cell clone phylogeny inference and refinement** in the main **Methods**) stayed at a certain value below $p$, indicating that the model would overfit as $p$ got beyond 10. We then picked the solution corresponding to the $p$ value that resulted in the highest inter-cell heterogeneity, so as to provide the highest granularity to ConDoR. Although this usually would still result in overfitting CN clones, a refinement step post-ConDoR would merge such clones that had no loss of heterozygosity (LOH) events, which can be reliably measured from targeted scDNA-seq data, between them. This ensured that the resulting copy number clones were supported by both total and allele-specific CNVs.

#### 2 Cancer tumor phylogeny reconstruction

We reconstruct the evolutionary history of the tumor under the constrained $k$-Dollo model. The constrained $k$-Dollo model integrates SNVs and SNPs with partial information about CNAs on the same set of cells during phylogeny inference. Specifically, the model incorporates CNAs via a clustering of cells, where all cells in the same cluster have the same copy-number profile (the copy number of all loci across the genome). The constrained $k$-Dollo model involves the following four constraints. First, a mutation is allowed to be gained at most once in the phylogeny. This constraint stems from the *infinite-sites assumption* which posits that it is very unlikely for the same position in the genome to get mutated multiple times independently. Second, a mutation can be lost at most $k$ times in the phylogeny, where $k$ is user-defined parameter. Third, since reversal of mutations (*back mutations*) in cancer are rare [1], we assume that SNVs and SNPs can only be lost due to overlapping CNAs. As such, mutation losses are only allowed between cells that belong to distinct copy-number clusters. Fourth, we assume that each copy-number profile describing the copy number states over the entire genome arises only once in the phylogeny. As a consequence, all cells belonging to the same copy number cluster form a connected subtree in the phylogeny.

Recently, Sashittal *et al.*, introduced ConDoR to infer tumor phylogenies at single-cell resolution from targeted single-cell sequencing data under the constrained $k$-Dollo model. ConDoR uses a probabilistic model of the observed read counts for each mutation to address errors and missing entries in the data. Here, we develop a scalable version of ConDoR to build coarse-grained tumor phylogenies at the copy number cluster level. Specifically, each mutation can have one of the following four states for each copy number cluster – (i) none of the cells contain the mutation (state $0$), (ii) some of the cells contain the mutation (state $(+)$), (iii) all the cells contain the mutation (state $1$), and (iv) all the cells have lost the mutation (state $2$). Under the constrained $k$-Dollo model, since a mutation can occur at most once in the phylogeny, each mutation can be gained in at most one of the copy number clusters. As such, for each mutation, there can be at most one copy number cluster with state $(+)$. Using the above model results in a phylogeny where each leaf is a copy-number clone and edges are annotated by gain or loss of mutations. This phylogeny is refined to generate a single-cell phylogeny using the procedure described in the following subsection.

##### 2.1 Subclonal SNV Phylogeny Reconstruction

For each copy number clone, we ConDoR [2] on these mutations to build a subclonal phylogeny. The resulting sub-clonal phylogeny is then used to refine the at the copy number clone leaf in the coarse phylogeny to construct the single-cell phylogeny. We describe the read count model in the next section.

#### 3 Read count model

We extended the mutation genotype model used by ConDoR [2] to include more genotypes: wild-type, heterozygous, homozygous, and deleted. We assume that measurement of mutation across cells is independent. As such Let $s_{i,\ell}$ denote the state of mutation $j$ in copy number cluster $\ell$. Let $q_{i,j}$ and $r_{i,j}$ denote the variant reads and total reads of mutation $j$ in cell $i$, respectively. Let $\sigma(i)$ denote the copy number cluster of cell $i$.

$$\Pr(Q \mid R, S) = \prod_{i=1}^{n}\prod_{j=1}^{m}\Pr(q_{i,j} \mid r_{i,j}, s_{i,j}).$$

Similar to previous papers [2, 3], we use the beta-binomial model for the read counts $r_{i,j}$ of each mutation in each cell. When the mutation is present, i.e. $s_{i,j} = 1$, we model the the read counts as follows,

$$q_{i,j} \mid s_{i,\sigma(i)} = 1 \sim \text{Beta-Binom}(r_{i,j}, \alpha, \beta),$$

where we set $\alpha = \beta = 1$ for this study. When the mutation is absent, i.e. $s_{i,j} = 0$, there may still be variant reads, i.e. $r_{i,j} > 0$, because of sequencing errors. Let $f_p$ be the sequencing error rate to produce a false positive variant read and $d$ be the dispersion parameter to account for allele dropouts. We model the read counts for $s_{i,j} = 0$ case as follows,

$$q_{i,j} \mid s_{i,\sigma(i)} \in \{0, 2\} \sim \text{Beta-Binom}(r_{i,j}, f_p d, (1 - f_p)d).$$

Finally, for the gain state, i.e. $s_{i,j} = (+)$, we do not impose presence of absence of mutations in the cells of the copy number cluster. We model this as follows

$$\Pr(q_{i,j} \mid s_{i,\sigma(i)} = (+)) = \max(\Pr(q_{i,j} \mid s_{i,\sigma(i)} = 0), \Pr(q_{i,j} \mid s_{i,\sigma(i)} = 1)).$$

In this study, we run ConDoR with a false-positive parameter $f_p = 0.001$ and dispersion parameter $d = 50$.

### 4 Mutual exclusivity

Suppose we have a probability matrix $P \in [0, 1]^{n \times m}$ for $n$ cells and $m$ mutations, where $p_{i,j}$ is the probability that mutation $j$ is present in cell $i$. For any two mutation $j$ and $j'$, let $Y$ be the random variable that denote the number of observations of mutual exclusivity across the $n$ cells, i.e. the number of observations of either $(0, 1)$ or $(1, 0)$ gametes in the mutation matrix. Assuming that the mutation states of cells are independent, $Y$ is a Poisson binomial variable with parameters

$$\mathbf{p} = \{p_{i,j}(1 - p_{i,j'}) + (1 - p_{i,j})p_{i,j'} : \forall i \in [n]\}.$$

We define a null distribution that conserves the marginal expectation of the number of occurrences of each mutation across the $n$ cells. Specifically, we define probability matrix $P'$ such that

$$p'_{i,j} = \frac{\sum_{i=1}^{n} p_{i,j}}{n}.$$

Let $X$ be the random variable that denote the number of observations of mutual exclusivity for mutations $j$ and $j'$ across the $n$ cells under the null distribution. $X$ is a binomial variable with $n$ trials and probability of success given by

$$p = \frac{\sum_{i=1}^{n} p_{i,j}}{n} \left(1 - \frac{\sum_{i=1}^{n} p_{i,j'}}{n}\right) + \left(1 - \frac{\sum_{i=1}^{n} p_{i,j}}{n}\right) \frac{\sum_{i=1}^{n} p_{i,j'}}{n}.$$

The p-value for rejecting the null hypothesis is given by $\Pr(X \geq Y)$. We formally define hypothesis testing problem as follows.

**Problem 4.1** (Mutual exclusivity test)**.** *Given a probability matrix $P$ and two mutation $j$ and $j'$, find the p-value $\Pr(Y \leq X)$, where $Y$ and $X$ is the number of occurrences of gametes $(0, 1)$ and $(1, 0)$ for mutation $j$ and $j'$ under the alternate and the null hypothesis, respectively.*

We compute the p-value $\Pr(Y \leq X)$ by marginalizing $X$ as follows,

$$\begin{aligned}\Pr(Y \leq X) &= \sum_{k=1}^{n} P(Y \leq k) \Pr(X = k) \\ &= \sum_{k=1}^{n} P(Y \leq k) \binom{n}{k} p^k (1-p)^{n-k},\end{aligned}$$

where $\Pr(Y \leq k)$ is the cumulative probability of the Poisson Binomial variable, computed using the poibin package [4]. In this study, we employ the read count model (Section 3) to determine the input probability matrix $P$.

### 5 ConDoR Parameter Tuning

We tuned the false-positive $f$ and ADO precision $a$ parameters of fast-ConDoR through a series of qualitative analyses of samples in our dataset. We selected a set of mutation of interest $m^*$ and manually annotated the expected state that fast-ConDoR should output for $m^*$ across all copy number clones with the read count heatmaps as a reference for our manual annotations. For multi-sample datasets, we provided manually annotated states across all samples. We then performed a grid search across varying values ($f \in [0.003, 0.03]$, $a \in [5, 91]$ ) for the two parameters and selected the set of parameters $f^*, a^*$ that maximizes the agreement between the fast-ConDoR results and the manual annotations.

### 6 Validating total copy number, LOH, homdel, amplification calling from FALCON and CONDOR

#### 6.1 Validation by colocalization of events

We validated LOH, homdel inferred by Tapestri-CN by identifying distinguishing somatic variants that can be reliably measured using Tapestri data in the same set of cells. Specifically, we observed several cases in which presence/absence of homdels coincided with LOH events and gain of somatic mutations. The likelihood of these observations by random chance is very low. In the following, we describe some of these cases.

1. In PC09, Tapestri-CN identifid *CDKN2A* homdel in cells that also have somatic mutations to driver genes such as *KRAS*, *TP53 ARID2*, and LOH in *STK1* and *WRN* 1a.
2. In PC12, Tapestri-CN identified *CDKN2A* homdel in cells that also have somatic mutations in *KRAS* and *TP53*. Specifically in this case, although *CDKN2A*'s amplicons 3, 4, 5 (spanning chr9:21970743-21974717) showed homdel signal in the normal clone (magenta, bottom

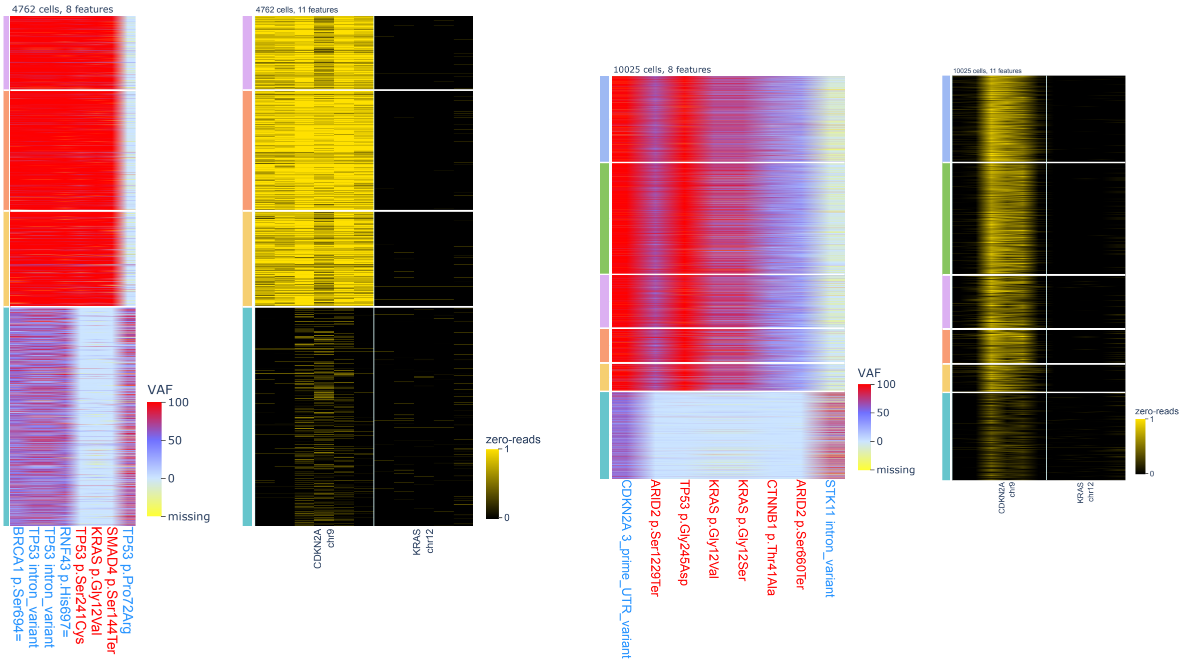


**(a)** **(b)**

**Supplementary Methods Figure 1: single-cell heatmaps** showing: VAf signal of select SNP/SNVs (left of each panel); homdel signal (per-amplicon read count = 0) of amplicons of select genes, for 1a: PC09. 1b: PC12. Each row is one single cell while each column is a feature (mutation or amplicon).

clone), they had significantly more prevalent signal in the cancer clones, indicating a likely homdel. Interestingly, amplicons 1, 2, 6 did not show homdel signal in the cancer clones, indicating that only part of the *CDKN2A* gene was deleted.

##### 6.2 validation by comparison to bulk CN calling

Orthogonally, we ran HATCHet2 on a subset of the Tapestri cases which had multiregional bulk whole-genome sequencing data (N=6). HATCHet2 is a state-of-the-art algorithm that increases the confidence of allele-specific copy number calling by leveraging signals from multiple samples of the same tumor [5], and we sought to compare its resulting total copy number clones, LOH/homdel/amplification events against those called by Tapestri.

###### 6.2.1 Tapestri inferred similar yet more granular clones than HATCHet2

Due to the large difference in the genomic resolution of whole genome sequencing and targeted single-cell DNA sequencing, we do not expect an exact match between the total copy number clones inferred by HATCHet2 and Tapestri-CN. Additionally, HATCHet2 is able to resolve whole-genome doubling (WGD) while Tapestri-CN cannot model that. Nevertheless, we observed general agreement between the clones inferred by the two methods.

For instance, in case PC06, HATCHet2 inferred clonal WGD, so for comparison of total copy number clones with Tapestri-CN we had to scale down every total copy number profile by 2. Then, we observed that both HATCHet2 and Tapestri-CN inferred clonal or near clonal loss at chromosome 2p (*DNMT3A, MSH2*), 6q (*ARID1B*), 9p and 9q (*CDKN2A, TGFBR1*), 10q (*BMPR1A*), 13q (*BRCA2)*, 17p (*TP53*), 18q *(SMAD2, SMAD7, SMAD4)*; amplification at chromosome 2q (*IDH1, BARD1*) (**Supplementary Methods Figure** 2a, 2b).

But first, Tapestri-CN inferred more tumor clones (n=4) than HATCHet2 (n=2). Next, Tapestri-CN inferred novel subclonal loss at chromosome 4q (*BMPR1B*), 11q (*ATM*); subclonal amplification at chromosome 16p (*CREBBP*). Importantly, Tapestri captured subclonal *CDKN2A* (chromosome 9p) deletion that HATCHet2 missed (**Supplementary Methods Figure** 2b).

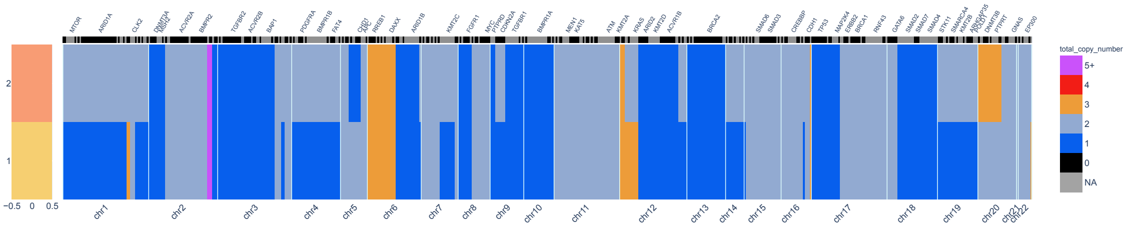


**(a)**

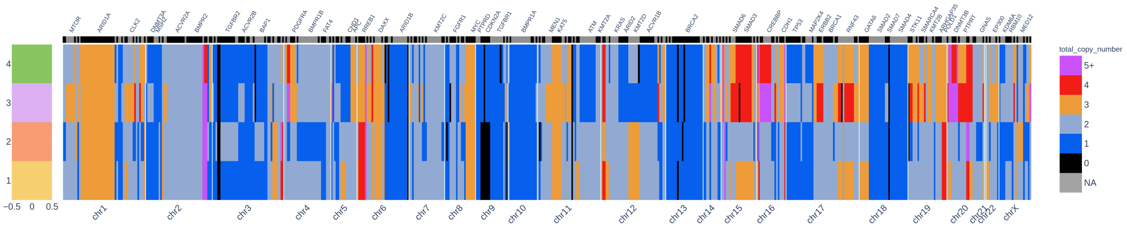


**(b)**

**Supplementary Methods Figure 2: PC06 Total copy number clones inferred by HATCHet2 compared with Tapestri-CN** (a) Total copy number clone profiles inferred by HATCHet2, subset to genomic locations covered by our Tapestri panel. Clone labels and colors on the left are arbitrary and do not necessarily map to those in the plot below. (b) Total copy number clone profiles inferred by Tapestri-CN.

Due to higher resolution of scDNA-seq, Tapestri-CN inferred generally higher number of copy number clones in every patient (**Supplementary Methods Figure** 3a). We then tried to match the nearest Tapestri-CN clone to each HATCHet2 clone and computed the error in the total copy number inferred for each of the 293 genes covered by the Tapestri panel. As expected, the mismatch in the inferred copy numbers is lower when the number of amplicons targeting a gene is higher (**Supplementary Methods Figure** 3b).

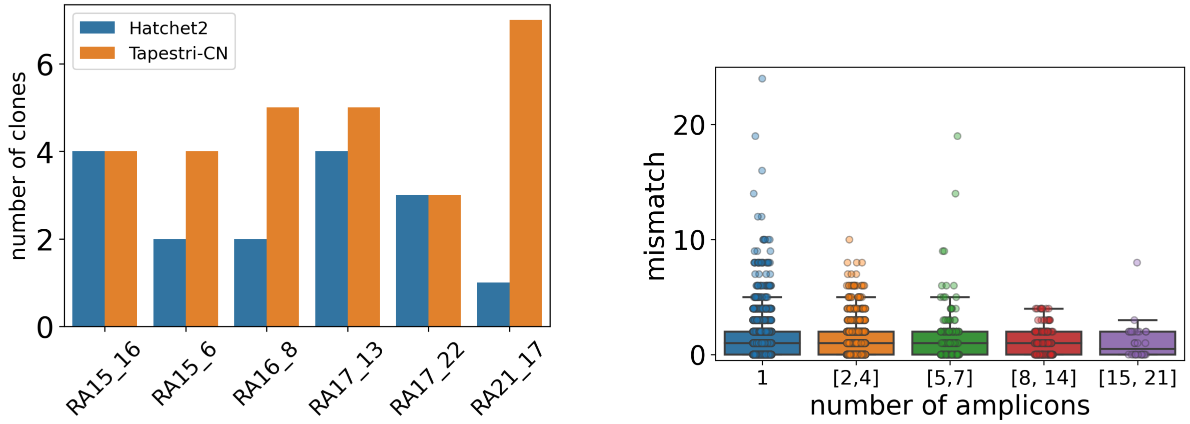


**(a)** **(b)**

**Supplementary Methods Figure 3: Copy number clones inferred by Tapestri-CN are consistent with Hatchet2 clones.** (a) Number of clones inferred by Hatchet2 and Tapestri-CN for 6 different patients. (b) Mismatch between the inferred copy number of genes in cancer cells between Hatchet2 and Tapestri-CN for increasing number of amplicons. Box plots show the median and the interquartile range (IQR), and the whiskers denote the lowest and highest values within 1.5 times the IQR from the first and third quartiles, respectively.

##### 6.2.2 Tapestri captured loss of heterozygosity events detected by HATCHet2

We sought to verify if select LOH events called by Tapestri from the mutation VAF signal were also detected by HATCHet2 which used SNP b-allele frequency (BAF) to detect LOH. For this purpose, we selected samples of high tumor purity to maximize the signal recovered from bulk, and plotted the BAF distribution calculated by HATCHet2 around genes called as LOH by Tapestri. As an example, in PC11 from the HATCHet2 side, we saw clear BAF signal from HATCHet2 indicating LOH at *KRAS, TP53* (**Supplementary Methods Figure** 4 **a, b**), which were both captured by Tapestri (**Supplementary Methods Figure** 4 **c, d**).

##### 6.2.3 Tapestri captured focal amplification events supported by HATCHet2

Next we wanted to see if focal amplification called by Tapestri was supported by HATCHet2.

As an example, as described in the main text, Tapestri detected amplifications to *MYC, MTOR, GATA6* in PC11 which were manifestation of continuous evolution in this tumor (**Supplementary Methods Figure** 5 **e**). In parallel, HATCHet2 showed BAF signals indicating allelic imbalance in these three genes (**Supplementary Methods Figure** 5 **a,b,c**), while the fraction copy number calls validated the high copy number state of *MYC* (**Supplementary Methods Figure** 5 **d, red circled)** which might be caused by extrachromosomal DNA as described in the main text.

##### 6.2.4 Tapestri captured focal homdel events with traces in HATCHet2

One of Tapestri-CN's key strength is detecting focal homdel as small as a single amplicon (200-300 base pairs). While HATCHet2 uses average genomic bin size of ∼100 kilobasepair (kb) and

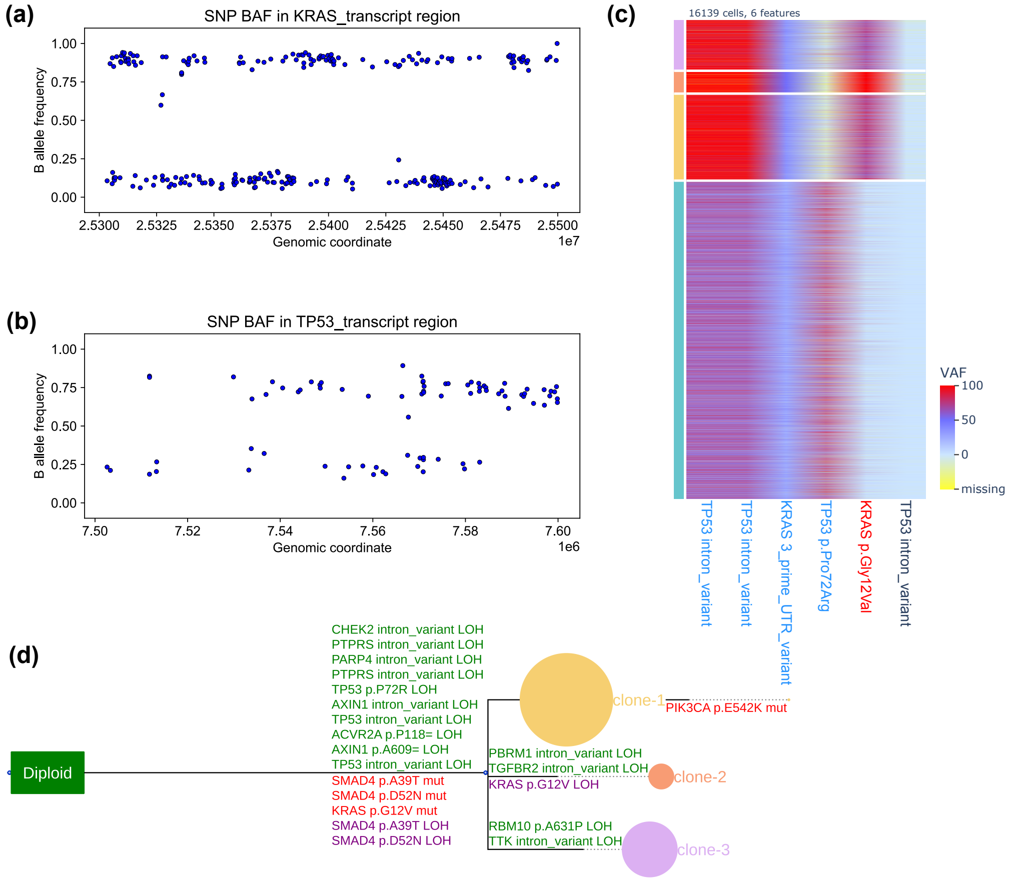


**Supplementary Methods Figure 4: PC11 HATCHet vs Tapestri-CN LOH comparison.** SNP BAF distributions at the *KRAS* **(a)** and *TP53* **(b)** genomic regions from WGS of the highest tumor purity sample of PC11. **(c)** Tapestri single-cell mutation VAF heatmap, split by ConDoR-inferred clones. Note the VAF shift in the tumor clones (pink, orange, yellow). **(d)** ConDoR-inferred phylogeny, with germline SNP LOH and somatic SNV gain/LOH labeled on the branches. Not the *KRAS, TP53* LOHs.

cannot capture such events, it did show some traces of them.

In case PC11, Tapestri captured an LOH of 1 of 11 amplicons of *SMAD4*, and homdel of the remaining 10 of 11 amplicons covering the gene (∼30kb) (**Supplementary Methods Figure 6 (a)**). On the other hand, HATCHet2 called LOH with bin size of 240kb (48.41 - 48.65 megabasepair, mb) encompassing most of the *SMAD4* gene. Nevertheless, while the flanking regions' BAFs suggested loss of heterozygosity (LOH), the region corresponding to the homdel called by Tapestri showed balanced B-allele frequencies (BAFs) with HATCHet2, likely suggesting homdel (48.575 - 48.625mb, **Supplementary Methods Figure 6 (b)**).

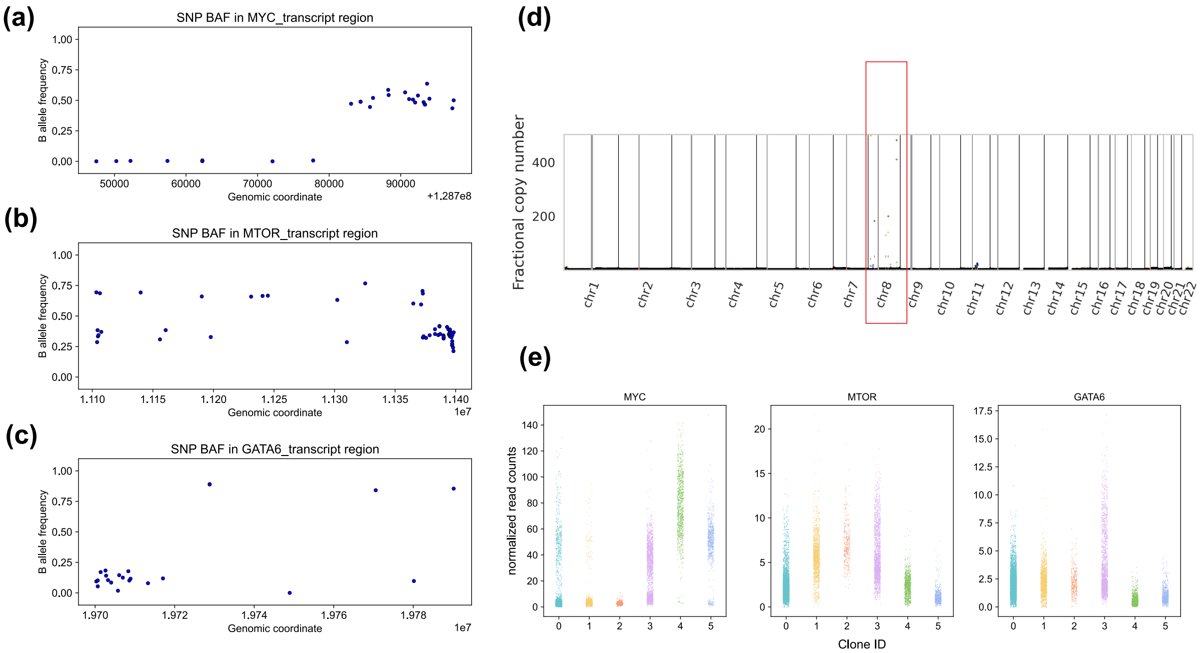


**Supplementary Methods Figure 5: PC10 HATCHet2 vs Tapestri-CN amplification comparison** SNP BAF distributions at the *MYC* (**a**), *MTOR* (**b**), *GATA6* (**c**) genomic regions from WGS of PC10's liver met which was also studied by Tapestri. **(d)** HATCHet2 genome-wide fractional copy number result. Note the high copy number state at *MYC* (chromosome 8, red circled). **(e)**, from Tapestri, normalized read count distributions for *MYC, MTOR, GATA6* per clone. Note the amplifications in the tumor clones.

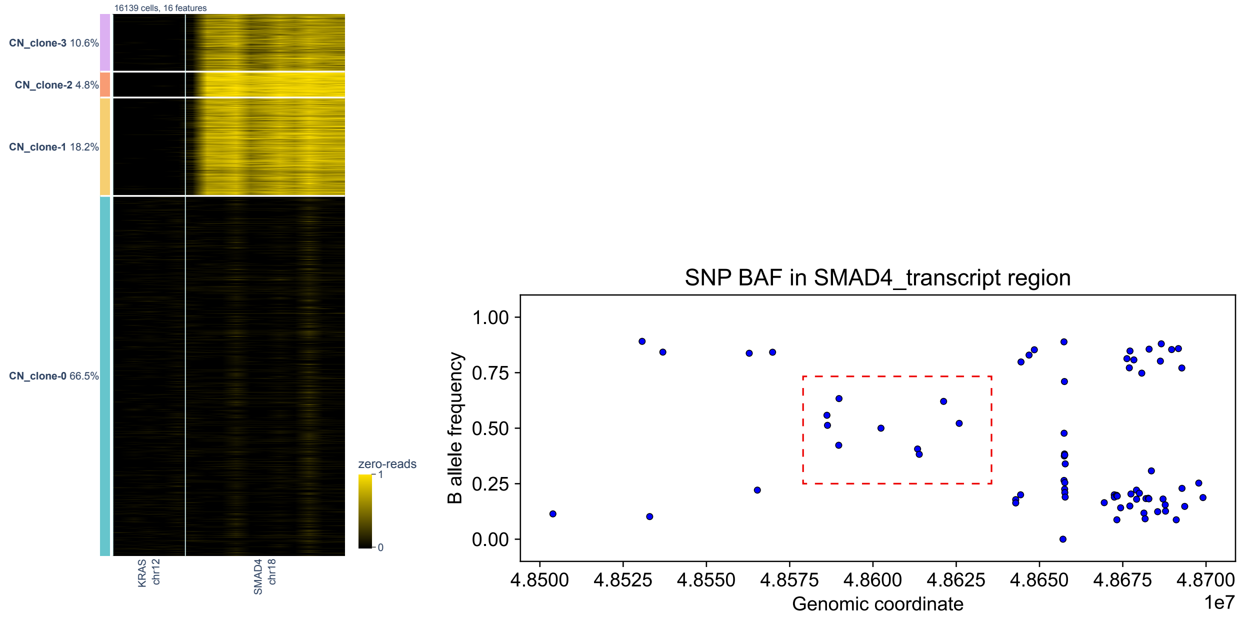


(a) (b)

**Supplementary Methods Figure 6: Tapestri detected a novel *SMAD4* focal homdel, with traces from HATCHet2**. **(a)** single-cell heatmap showing homdel signals of genes of interest. Note the homdel of 10 of 11 amplicons targeting *SMAD4*. **(b)** SNP BAF distribution at the *SMAD4* genomic region from WGS of PC11's liver met which was also studied by Tapestri. Note the balanced BAF region (red circled) flanked by the unbalanced BAF regions
